# Combinatorial drug-microenvironment interaction mapping reveals cell-extrinsic drug resistance mechanisms and clinically relevant patient subgroups in CLL

**DOI:** 10.1101/2021.07.23.453514

**Authors:** Peter-Martin Bruch, Holly A. R. Giles, Carolin Kolb, Sophie A. Herbst, Tina Becirovic, Tobias Roider, Junyan Lu, Sebastian Scheinost, Lena Wagner, Jennifer Huellein, Ivan Berest, Mark Kriegsmann, Katharina Kriegsmann, Christiane Zgorzelski, Peter Dreger, Judith B. Zaugg, Carsten Müller-Tidow, Thorsten Zenz, Wolfgang Huber, Sascha Dietrich

## Abstract

The tumour microenvironment and genetic alterations collectively influence drug efficacy in cancer, but current evidence is limited to small scale studies and systematic analyses are lacking. We chose Chronic Lymphocytic Leukaemia (CLL), the most common leukaemia in adults, as a model disease to study this complex interplay systematically. We performed a combinatorial assay using 12 drugs individually co-applied with each of 17 microenvironmental stimuli in 192 primary CLL samples, generating a comprehensive map of drug-microenvironment interactions in CLL. This data was combined with whole-exome sequencing, DNA-methylation, RNA-sequencing and copy number variant annotation. Our assay identified four distinct CLL subgroups that differed in their responses to the panel of microenvironmental stimuli. These subgroups were characterized by distinct clinical outcomes independently of known prognostic markers. We investigated the effect of CLL-specific recurrent genetic alterations on microenvironmental responses and identified trisomy 12 as an amplifier of multiple microenvironmental stimuli. We further quantified the impact of microenvironmental stimuli on drug response, confirmed known interactions such as Interleukin (IL) 4 mediated resistance to B cell receptor (BCR) inhibitors, and identified new interactions such as Interferon-γ induced resistance to BCR inhibitors. Finally, we identified interactions which were limited to genetic subgroups. Resistance to chemotherapeutics, such as Fludarabine, induced by Toll-Like Receptor (TLR) agonists could be observed in IGHV unmutated patient samples and IGHV mutated samples with trisomy 12. In-vivo relevance was investigated in CLL-infiltrated lymph nodes, which showed increased IL4 and TLR signalling activity compared to healthy samples (p<0.001). High IL4 activity in lymph nodes correlated with faster disease progression (p=0.038).

We provide a publicly available resource (www.dietrichlab.de/CLL_Microenvironment/) which uncovers tumour cell extrinsic influences on drug response and disease progression in CLL, and how these interactions are modulated by cell intrinsic molecular features.

## Introduction

CLL is a common B-cell malignancy, attributed to the accumulation of mature B-lymphocytes in the peripheral blood, bone marrow, and lymph nodes^1^. The disease course is heterogeneous, influenced by multiple factors including B cell receptor (BCR) signalling, genetic alterations, epigenetic effects, and the tumour microenvironment^2^. Despite significant improvements to CLL treatment, including the advent of BCR inhibitors^3^ and BH3-mimetics^4^, CLL remains incurable. There is a compelling need to better understand cell-intrinsic and extrinsic causes of therapy failure.

The genetic landscape of CLL is well characterised. The most important features include del(17p), TP53-mutations, immunoglobulin heavy chain gene (IGHV) mutation status, del(13q) and trisomy 12^5, 6^. This list of recurrent genetic and structural alterations has expanded in the past years^7^. The predictive value of those alterations has however declined in the context of modern targeted therapies and resistance mechanisms are incompletely understood ^8^.

Additionally to cell-intrinsic factors, CLL cell proliferation and survival is dependent on the lymph node microenvironment^9^, which is underlined by the observation that CLL cells undergo spontaneous apoptosis in vitro if deprived of protective microenvironmental signals^10^. Microenvironmental stimuli can induce drug resistance in vitro, for example the combination of IL2 and resiquimod, a Toll-Like Receptor (TLR) 7 / 8 stimulus, induces resistance to venetoclax^11^. The CLL microenvironment constitutes a complex network of stromal and immune cells that promote cell expansion^12^ via soluble factors and cell-cell contacts. Microenvironmental signalling is particularly important in protective niches, especially lymph nodes. Incomplete response to BCR inhibitors has been linked to persistent enlargement of lymph nodes^13^, which are the main site of CLL proliferation^9^.

Several studies have investigated individual components of the microenvironment in leukaemia^14–18^. However, systematic studies, particularly those exploring cell-extrinsic influences on drug response, are rare. Carey et al.^19^ have screened acute myeloid leukaemia samples with a panel of soluble factors and among them identified IL1ß as a mediator of cellular expansion in Acute Myeloid Leukaemia. This example highlights the value of systematic approaches to dissect tumour-microenvironment crosstalk.

Taking this approach further, we screened a panel of microenvironmental stimuli in CLL, individually and in combination with drugs, and complemented our dataset with multi-omics data on the patient samples. In this study, we integrate genetic, epigenetic and microenvironmental modulators of drug response in CLL systematically in a large patient cohort that covers the clinical and molecular diversity of CLL. Due to the dependency on microenvironmental support, CLL is an important model system for the interaction between malignant and microenvironmental cells in general.

## Methods

### Sample preparation and drug-stimulation profiling

Sample preparation, cell-culture, drug-stimulation profiling, and genomic annotation was performed on 192 CLL patient samples as previously described^20^ with the following adjustments. Stimuli and drugs were mixed and preplated in the culture plates directly before adding the cell suspensions. RPMI-1640 and supplements were acquired from Gibco by Life Technologies, human serum was acquired from PAN Biotech (Cat.No. P40-2701, Lot.No:P- 020317). Viability was assessed by measuring ATP concentration using CellTiter-Glo (Promega) after 48h. Luminescence was measured on a Perkin Elmer EnVision.

### Compounds and Stimuli

Compounds and stimulatory agents were dissolved, stored, and diluted according to manufacturer’s protocol. HS-5 conditioned medium was produced by incubating HS-5 stromal cell line to >80% confluency and cell removal by centrifugation. For a detailed list of stimuli and drugs and associated concentrations, see Supp. Tables 1 and 2. Final DMSO concentration did not exceed 0.3%.

### Immunohistochemistry

Lymph node biopsies of CLL-infiltrated and non-neoplastic samples were formalin fixed, paraffin embedded, arranged in Tissue Microarrays and stained for pSTAT6 (ab28829, Abcam) and pIRAK4 (ab216513, Abcam). The slides were analysed using Qupath^21^ and the recommended protocol.

### Data processing and statistical analysis

To quantify the responses to drugs and stimuli, we used a measure of viability relative to the control, namely the natural logarithm of the ratio between CellTiter Glo luminescence readout of the respective treatment and the median of luminescence readouts of the DMSO control wells on the same plate, excluding controls on the outer plate edges. Data analysis was performed using R version 4 and using packages including DESeq2^22^, survival^23^, Glmnet^24^, ConsensusClusterPlus^25^, clusterProfiler^26^, ChIPseeker^27^, genomation^28^ and BloodCancerMultiOmics2017^29^ to perform univariate association tests, multivariate regression with and without lasso penalization, Cox regression, generalised linear modelling and clustering. The complete analysis is described along with computer-executable transcripts at github.com/Huber-group-EMBL/CLLCytokineScreen2021.

For Figure 2A, Clusters were determined by repeated hierarchical clustering over 10.000 repetitions with randomly selected sample subsets of 80% with the Euclidean metric using ConsensusClusterPlus^25^. LDT was calculated as previously described^30^. For figure 2D Multinomial regression with lasso penalisation with a matrix of genetic features (p=39), and IGHV status (encoded as M = 1 and U = 0) was used to identify multivariate predictors of cluster assignment. Coefficients shown are mean coefficients from 50 bootstrapped repeats and error bars represent the mean ± standard deviation. Genetic features with >20% missing values were excluded, and only patients with complete annotation were included in the model (n=137).

**Figure 1.**
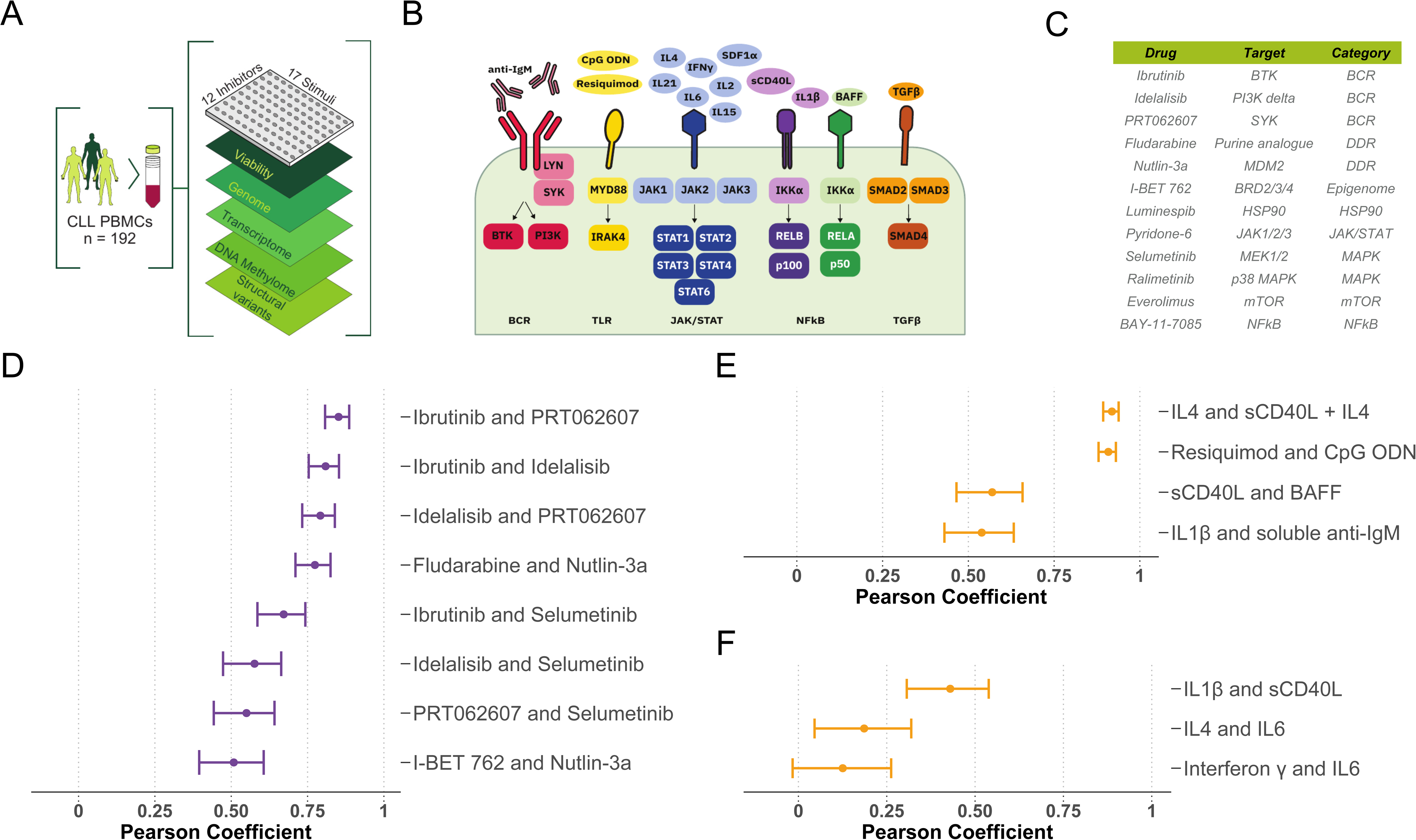
Study outline and overview of drugs and stimuli. (A) Schematic of experimental protocol. By combining 12 drugs and 17 stimuli, we systematically queried the effects of simultaneous stimulation and inhibition of critical pathways in CLL (n=192). Integrating functional drug-stimulus response profiling with four additional omics layers, we identified pro-survival pathways, underlying molecular modulators of drug and microenvironment responses, and drug-stimulus interactions in CLL. All screening data can be explored either manually (github.com/Huber-group-EMBL/CLLCytokineScreen2021) or interactively, via the shiny app (www.dietrichlab.de/CLL_Microenvironment/). (B) Overview of stimuli included in the screen and summary of their associated targets. HS-5 conditioned medium is omitted, as no specific target can be shown. (C) Table of drugs included in the screen. Drug target, and category of target are also shown. (D - F) Pearson correlation coefficients of drug-drug and stimulus-stimulus correlations. (D) and (E) show highest drug -drug and stimulus-stimulus correlations, (F) shows selected combinations of stimuli targeting the NfKB and JAK-STAT pathways as examples of stimuli which share very similar receptors and downstream target profiles but show a divergence of effects.

**Figure 2.**
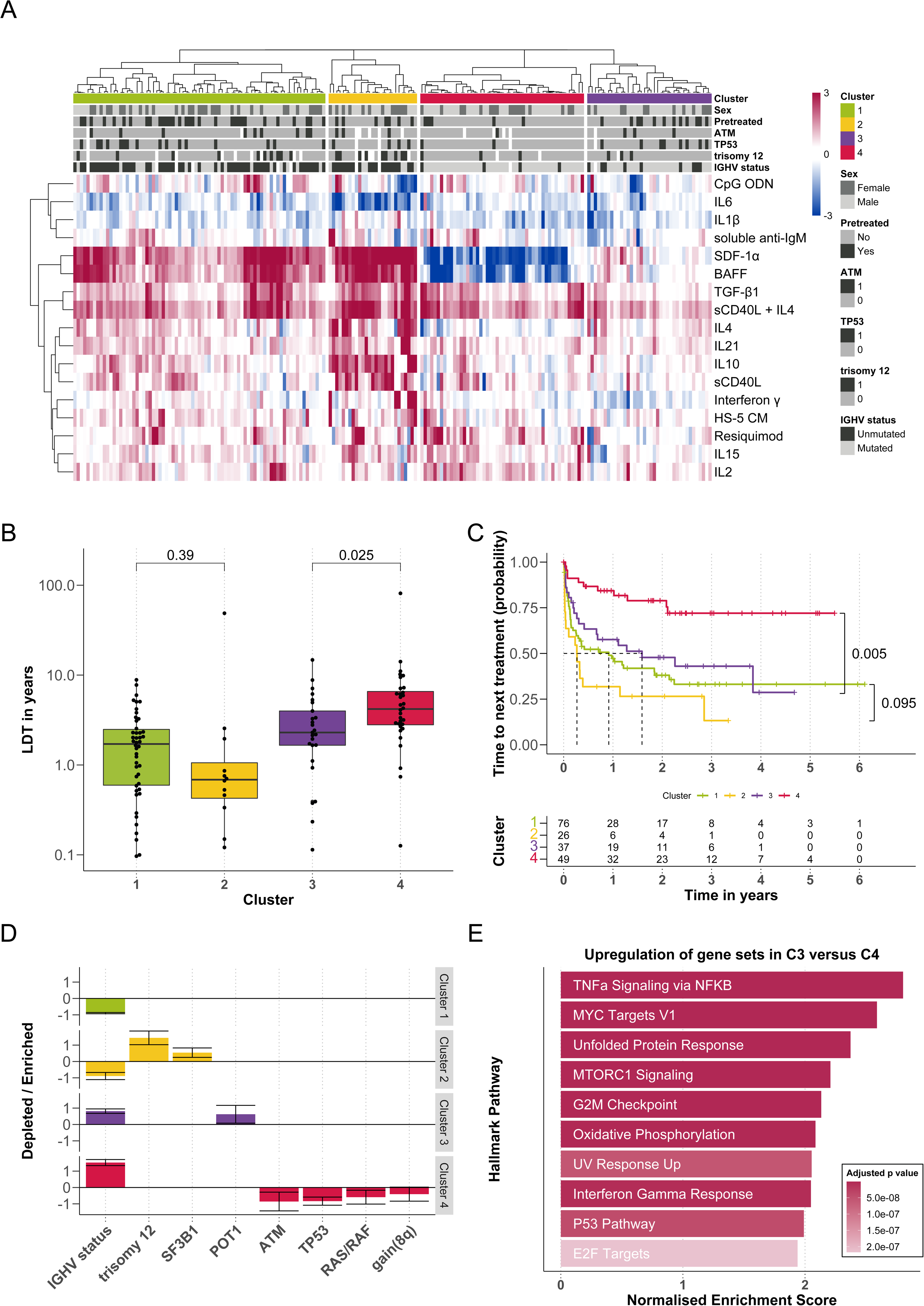
Ex vivo microenvironmental response profiling reveals subgroups with distinct molecular profiles and disease progression. (A) Heatmap showing the viability after treatment with microenvironmental stimuli. Rows represent microenvironmental stimuli and columns represent primary CLL samples, annotated for their genetic background, sex and pre-treatment status above. Red values indicate increased viability upon treatment, blue indicates decreased viability. Data is row-scaled. (B) Lymphocyte doubling time, a clinical marker for disease progression, plotted for each cluster. P-values from Student’s t-tests. (C) Kaplan-Meier curves of time to next treatment for each cluster. P-values from univariate Cox proportional hazard models comparing IGHV-U enriched C1 with C2, and IGHV-M enriched C3 with C4. (D) Multinomial regression with lasso penalisation to identify enrichment or depletion of genetic features within each cluster. (E) Gene set enrichment analysis (GSEA) comparing on expression of genes in samples from C3 and C4. Normalised enrichment scores are shown for top 10 most significant pathways upregulated in C3 versus C4.

Significance testing for genetic determinants of microenvironmental response shown in Figure 3A was performed for somatic mutations and copy number aberrations present in ≥3 patients, and IGHV status (n = 54). For multivariate modelling with L1 penalisation shown in Figure 3B, to generate the feature matrix genetic alterations with less than 20% missing values were considered, KRAS, BRAF and NRAS mutations were summarised as RAS/RAF alterations and only patient samples with complete genetic annotation were tested. In total, 39 genetic features as well as IGHV status (encoded as M = 1 and U = 0), and Methylation Cluster (encoded as 0, 0.5, 1) and 129 patients were included in this analysis.

**Figure 3.**
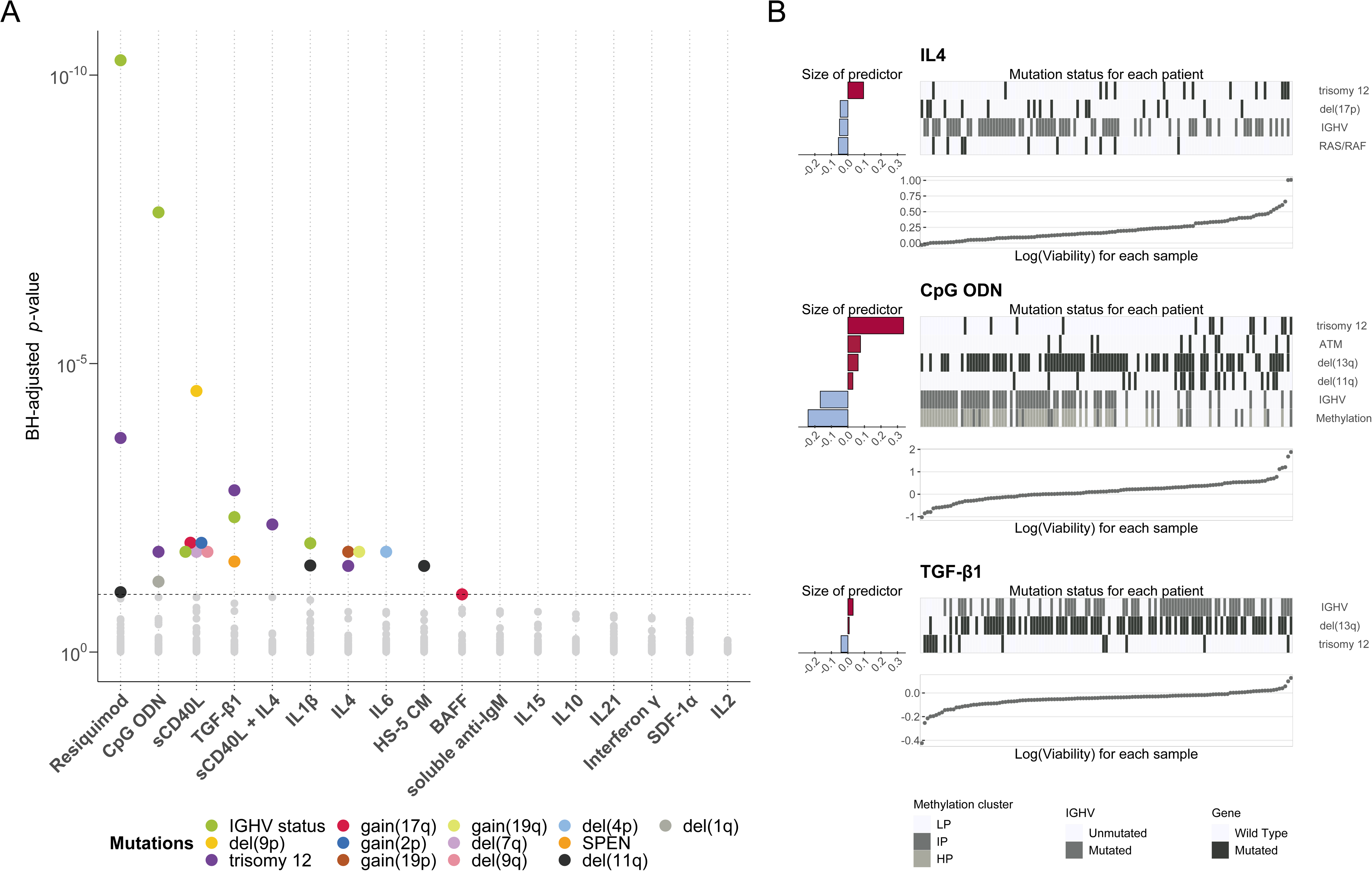
Responses to microenvironmental signals are modulated by genetic alterations recurrent in CLL. (A) Overview of results of all tested gene - stimulus associations. x-axis shows stimuli, y-axis shows p-values from Student’s t-test (two-sided, with equal variance). Each dot represents a gene-stimulus association. Tests with p-values smaller than the threshold corresponding to a false discovery rate (FDR) of 10% (method of Benjamini and Hochberg) are indicated by coloured circles, where the colours represent the gene mutations and structural aberrations. (B) Predictor profiles for selected stimuli to represent gene - stimulus associations identified through Gaussian linear modelling with L1-penalty. Within the plots, the bar plots on the left indicate size and sign of coefficients assigned to the named predictors. Positive coefficients indicate higher viability after stimulation if the feature is present. Scatter plots indicate log(viability) values, in order of magnitude, for each individual sample. Heatmaps show mutation status for each of the genetic predictors for the corresponding samples in the scatter plot.

For Figures 4C and D RNA-Seq data on matched samples belonging to clusters C3 and C4 were selected, Ig Genes were filtered out, differentially expressed genes were calculated with the design formula ∼ IGHV + Cluster using the Deseq2 package and ranked based on Wald statistics. GSEA was performed using the fgsea algorithm. For Fig 4E, we selected the raw sequencing files from Rendeiro et al. 2016^31^ to include one sample per patient passing quality checks (n=52). We obtained bam files mapped to the hg19 genome and adjusted for CG bias as previously outlined^32^. As trisomy 12 status was not already annotated, we called trisomy 12 in samples that contained > 1.4 times more reads per peak on average in chromosome 12, compared to peaks on other chromosomes. We used diffTf in permutation mode^32^ to infer the differential TF activity between trisomy 12 and non-trisomy 12 samples using HOCOMOCO v10^33^ with design formula: “∼ sample_processing_batch + sex + IGHV status + trisomy 12”. For Figures 4E and H, change in TF activity is inferred using diffTF^32^, and measured as weighted mean difference. For Figure 4F, we used ChIPseq data from Care et al. 2014^34^ to define TF targets as the closest gene to each significant ChIP peak (q value<0.05) and within ±1kb of Transcription Start Site. Over-representation tests were run using clusterProfiler package^26^ and the method corresponds to one-sided version of Fisher’s exact test. For Figure 4H, CLL cells have been treated for 6h hours with IBET-762 (1µM) or DMSO as solvent control before isolation and transposition of DNA for ATAC Seq. Raw ATACseq data generated from IBET-762 and DMSO treated CLL samples were processed as described in Berest et al. 2019^32^, mapped to hg38. We then used analytical mode of diffTF with HOCOMOCO v11 database^33^ using the following parameters: minOverlap = 2; design formula = “∼Patient + treatment. The plot depicts weighted mean difference of different diffTF analyses (trisomy 12 vs. non-trisomy 12 and IBET-762 treated cells vs. DMSO treated cells), absolute effect sizes should therefore not be directly compared. Significantly different TFs from the trisomy 12 vs. non-trisomy 12 analysis are shown.

**Figure 4.**
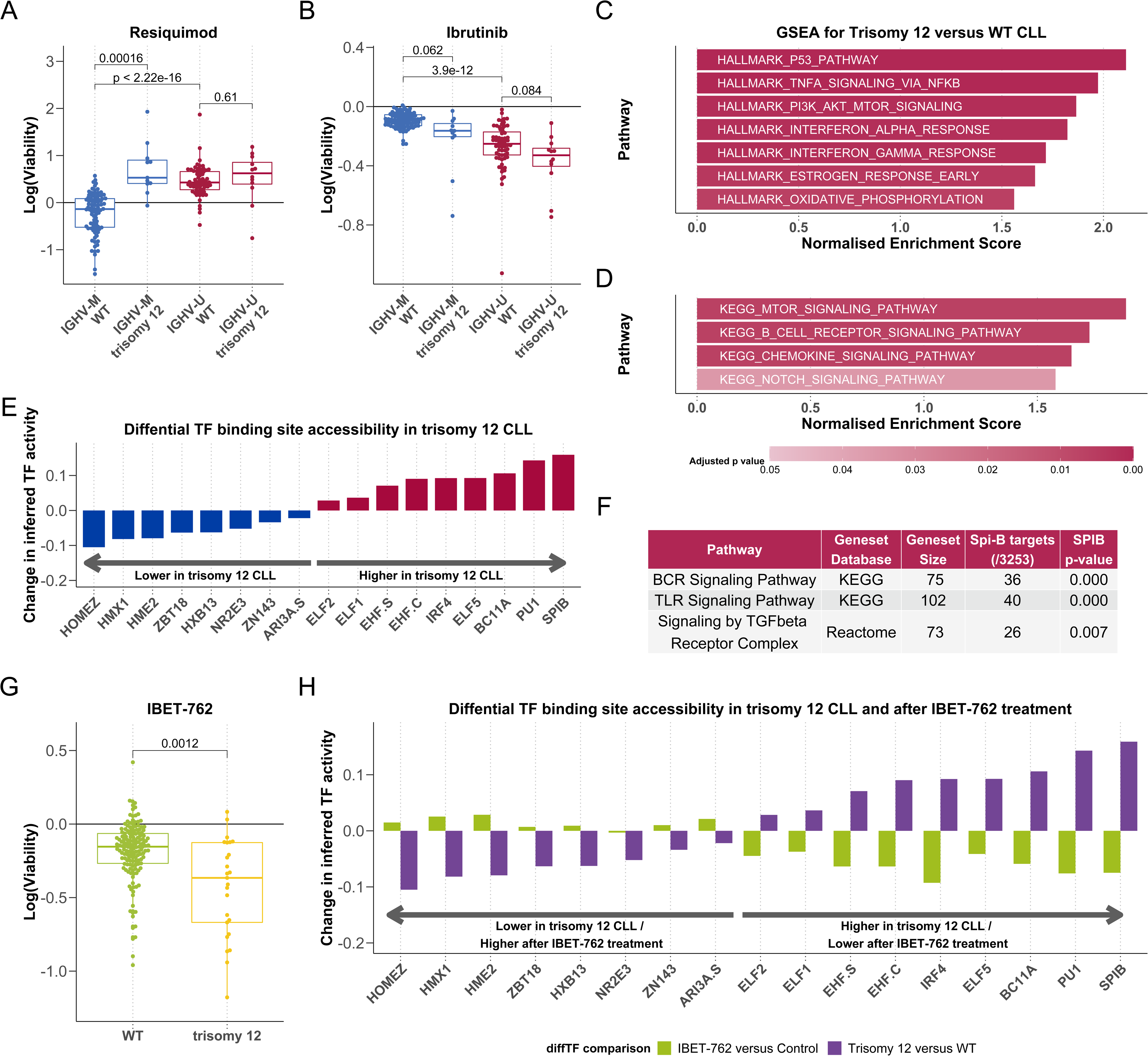
Trisomy 12 modulates responses to microenvironmental signals. (A+B) Log normalised viability after treatment with resiquimod (TLR 7/8 stimulus) (A) and ibrutinib (B) in the genetic subgroups of IGHV status and trisomy 12. P-values from Student’s t-tests. (C+D) GSEA comparing trisomy 12 samples to non-trisomy 12 samples in Hallmark genesets (C) and selected KEGG genesets of microenvironmental signalling pathways (D). (E) Differential TF accessibility (y axis) between trisomy 12 (n = 9) and non-trisomy 12 (n = 43) samples, data taken from Rendeiro et al^32^. TFs with adjusted p-value <0.05 are shown and ordered by change in inferred TF activity (x-axis). Spi-B and PU.1 TF binding sites show higher accessibility in trisomy 12 CLL. (F) Over-representation tests of selected KEGG and Reactome pathways in ChIPseq analysis of Spi-B binding in lymphoma cell lines indicates involvement in coordinating transcriptional response to microenvironmental signals, including BCR and TLR signalling. (G) Log normalised viability after treatment with the bromodomain inhibitor IBET-762 in trisomy 12 and non-trisomy 12 samples. P-value from Student’s t-tests. (H) Differential TF binding site accessibility (y axis) in trisomy 12 vs non-trisomy 12 CLL samples (purple) and for IBET-762 vs DMSO treated CLL samples (green). Direction of differential accessibility values are shown for two independent datasets comparing trisomy 12 vs non-trisomy 12 CLL and IBET-762 vs control-treated CLL, for all TFs with adjusted p value <0.05 in the trisomy 12 comparison. Absolute change in TF accessibility can not be compared between the two experiments.

To generate predictor profiles for Figure 6A, linear model in Eqn. (1) was fitted in a sample - specific manner, to calculate drug stimulus interaction coefficients (βint) for each patient sample. Associations between the size of βint and genetic features were identified using multivariate regression with L1 (lasso) regularisation with gene mutations (p = 39) and IGHV status as predictors and selecting coefficients that were chosen in >90% of bootstrapped model fits.

For Figure 7G+H pSTAT6 and pIRAK4 groups were defined using maximally selected rank statistics based on staining intensities. The same 64 CLL lymph node samples, for which survival data was available, were used for both Kaplan-Meier plots.

### Data Sharing Statement

Screening data and patient annotation used in this study are available at github.com/Huber-group-EMBL/CLLCytokineScreen2021. For original sequencing data and raw viability data please contact the corresponding authors. We obtained CLL ATACseq data^31^ from the EGA (EGAD00001002110) and ChIPseq data for Spi-B binding^34^ from the NCBI GEO database^35^ (GSE56857, GSM1370276).

## Results

### Ex-vivo cell viability assay demonstrates functional diversity of cytokine and microenvironmental signalling pathways

We measured the effects of 17 cytokines and microenvironmental stimuli on cell viability in 192 primary CLL samples and combined each with 12 drugs to investigate the influence on spontaneous and drug-induced apoptosis. (Fig 1A-C, Supp. Tables 1 -3). Viability was assessed by ATP measurement after 48h and normalised to untreated controls^20^.

The patient samples were characterised by DNA sequencing, mapping of copy number variants, genome-wide DNA-methylation profiling, and RNA-Sequencing^20^. The distribution of genetic features in our cohort is comparable to other studies (Supp. Fig 1)^6^.

To assess the heterogeneity of the response patterns, we calculated Pearson correlation coefficients for each pair of drugs and each pair of stimuli. For the drugs, high correlation coefficients (>0.75) were associated with identical target pathways. For example, drugs targeting the BCR pathway (ibrutinib, idelalisib, PRT062607 and selumetinib) were highly correlated, indicating that our data sensitively and specifically reflect inter-individual differences in pathway dependencies (Fig. 1D)^20^.

In contrast, microenvironmental stimuli showed lower correlations, even where stimuli targeted similar pathways (Fig. 1E, Supp. Fig. 2). For example, lower correlations were observed between different stimuli of the JAK-STAT and NfκB pathways, indicating a low degree of redundancy between stimuli. High correlations (R>0.75) were only seen with the TLR stimuli resiquimod (TLR7/8) and CpG ODN (TLR9) as well as IL4 and IL4 + soluble CD40L (sCD40L), which target near identical receptors and downstream targets.

Most stimuli increased CLL viability, underlining the supportive nature of the microenvironment. However, IL6, tumor growth factor β (TGFβ), and TLR7/8/9 agonists in IGHV-mutated (IGHV-M) samples decreased viability (Supp. Fig. 3). The strongest responses were seen with IL4 and TLR7/8/9 agonists, highlighting the potency of these pathways in modulating CLL survival. All screening data can be explored manually (github.com/Huber-group-EMBL/CLLCytokineScreen2021) and interactively, via the shiny app (www.dietrichlab.de/CLL_Microenvironment/).

### Patterns of responses to microenvironmental stimuli reveal two subgroups of IGHV-M CLL with differential aggressiveness

To gain a global overview over the pattern of responses to our set of microenvironmental stimuli across patient samples, we clustered^25^ and visualised the response profiles (Fig. 2A).

This analysis revealed four functionally defined subgroups with distinct response profiles. Two clusters (termed C1 and C2) were enriched in IGHV-Unmutated (IGHV-U), and two in IGHV- M CLL (C3 and C4). C1 and C2 showed strong responses to IL4 and TLR7/8/9 agonists. C2 was distinguished by stronger responses to the stimuli overall, in particular to NFκB agonists including IL1β and anti-IgM. C3 responded weakly to the majority of stimuli, while C4 was defined by a decreased viability upon TLR7/8/9 stimulation (Supp Fig. 4).

Next, we investigated whether these clusters were associated with differential in vivo disease progression of the corresponding patients. We used lymphocyte doubling time and time to next treatment to quantify the proliferative capacity of CLL cells. In line with clinical observations^36^, the IGHV-U enriched C1 and C2 showed a shorter lymphocyte doubling time than the IGHV-M enriched C3 and C4. More strikingly, C3 showed a significantly shorter lymphocyte doubling time than C4, indicating a higher proliferative capacity (Fig. 2B).

This observation was reflected in disease progression rates: The predominantly IGHV-M C3 showed a progression dynamic similar to the IGHV-U enriched C1 and C2. C4, however showed a longer time to next treatment than C1, C2 and C3 (Cox proportional hazards model, p= 0.0014, <0.001, 0.005 respectively, Fig. 2C, Supp. Fig 5A&B). This points towards a subset of mostly IGHV-M patients with more aggressive disease, which is characterised by its differential response to microenvironmental stimuli.

**Figure 5.**
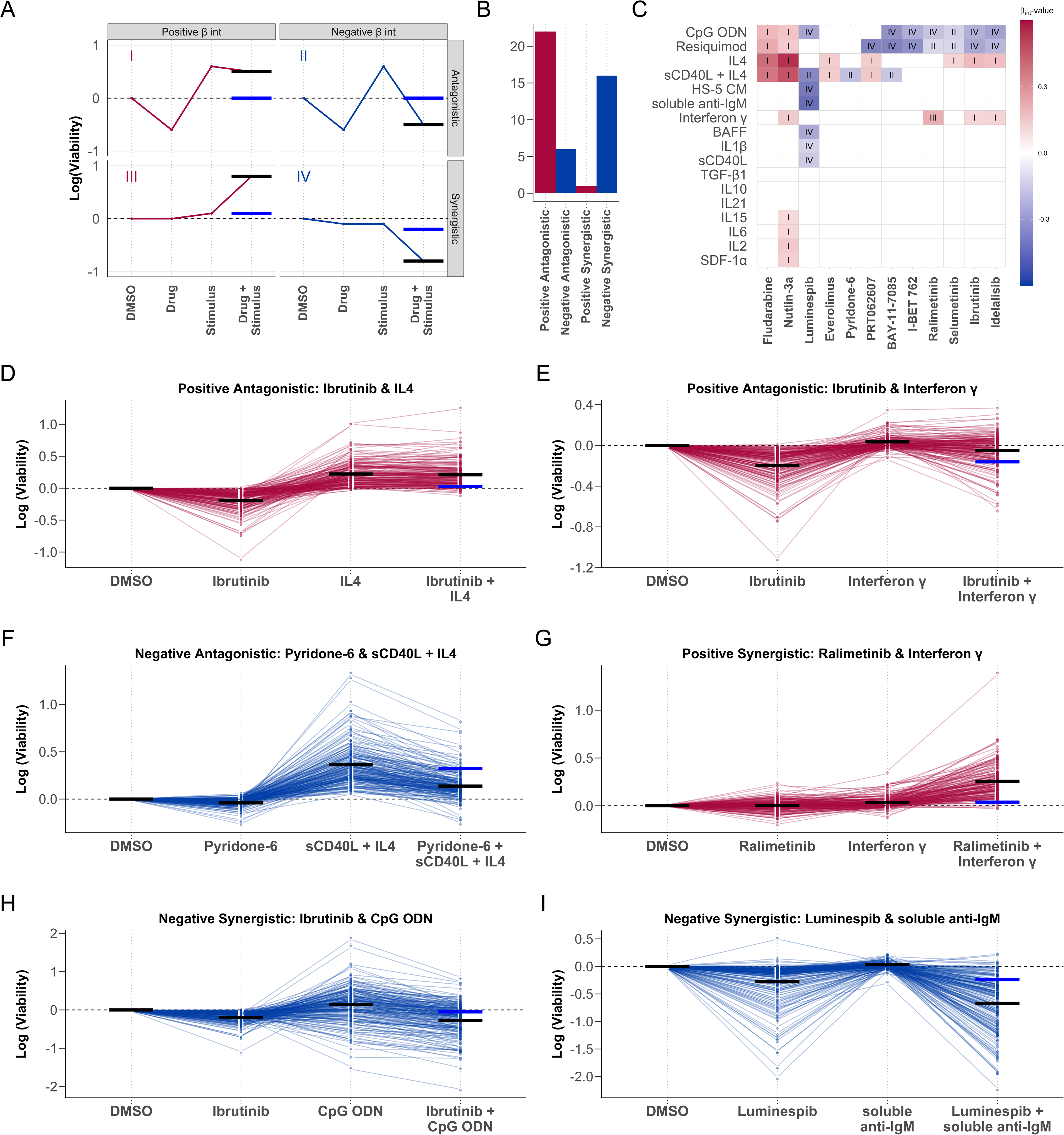
Microenvironmental stimuli influence ex-vivo drug response. (A) Graphical representation of the four drug-stimulus interaction categories. Categories are defined according to the nature of the interaction (synergistic or antagonistic), and whether the viability is increased or decreased by the stimulus (positive or negative). x-axis shows treatment type, y-axis shows viability with each treatment. Red and blue points and lines depict a representative treatment response pattern for given interaction type. Blue horizontal lines represent the expected viability for combinatorial treatment in the absence of an interaction between drug-stimulus (i.e. additive effects), black horizontal lines represent the measured viability after combinatorial treatment. The difference between the black and blue lines represents the drug - stimulus interaction. (B) Bar plot of significant interactions in all four categories (p-value for *β_int_* is <0.05). (C) Heatmap of all βint values for which p-value <0.05 for drug-stimulus combinations, annotated with interaction type. Scale indicates size and sign of βint. Rows and columns clustered according to hierarchical clustering. I: Positive *β_int_* and antagonistic (Microenvironmental stimulation reduces drug effect) II: Negative *β_int_* and antagonistic (Drug reduces stimuli effect) III: Positive *β_int_* and synergistic (Microenvironmental stimulation and drug have synergistic pro-survival effect) IV: Negative *β_int_* and antagonistic (Microenvironmental stimulation and drug show synergistic toxicity) (D - I) Examples of drug-stimulus interactions, for each category. Plots show log transformed viability values with each treatment, for all samples. Each line represents one patient sample linked across treatments. Black lines in single treatments indicate viability predicted by the linear model. In combinatorial treatment, the expected viability based on the additive effect of drug and stimulus (blue), and the viability with interaction (black), are shown to indicate the impact of the interaction. (D+E) Ibrutinib, a clinically used BTK inhibitor, is blocked by IL4 and IFNγ. (F) The JAK inhibitor pyridone-6 inhibits the pro-survival effect of sCD40L + IL4 stimulation. (G) The p38 inhibitor ralimetinib and IFNγ show a synergistic pro-survival effect not seen in either single treatment. (H) TLR agonists, including CpG ODN (shown) increase sensitivity to BTK inhibition by ibrutinib, despite increasing viability as single treatments. (I) Soluble anti-IgM sensitises CLL samples to HSP90 inhibition by luminespib.

To characterise genetic differences between clusters, we calculated predictor profiles for cluster assignment, using multinomial regression with the sample mutation profiles (Fig. 2D). IGHV status was the main predictor of all clusters. Trisomy 12 and SF3B1 mutations were associated with C2, which showed enhanced responses to many stimuli. C4, which was associated with slow in vivo progression, showed depletion of TP53, ATM, RAS/RAF mutations and gain(8q).

The observed difference in progression dynamics in vivo was not explained solely by the known prognostic markers IGHV, trisomy 12, and TP53. A multivariate Cox proportional hazards model accounting for these features showed an independent prognostic value of the cluster assignment between C3 and C4 (p=0.039, Supp. Table 4).

To examine why C3 exhibited faster disease progression than C4, we compared baseline pathway activity at the time of sampling, using RNA-Sequencing (Supp. Fig. 6A). Gene set enrichment analysis revealed that key cytokine gene sets were upregulated in C3, including TNFα signalling via NFκB, indicating higher pathway activity in vivo (Fig. 2E, Supp. Fig. 6B). In addition, pathways that indicate more aggressive disease relating to proliferation, metabolism and stress response were upregulated in C3 (Supp. Fig. 6 C-E).

Taken together, the patterns of responses to our set of stimuli distinguish two subgroups of IGHV-M CLL (C3 and C4) that are characterised by distinct in vivo pathway activities and differential disease progression.

### Microenvironmental signals and genetic features collectively determine malignant cell survival

Having observed heterogeneous responses to stimulation, we performed univariate analyses of genetic determinants of stimulus response, including IGHV status, somatic gene mutations, and structural variants (Fig. 3A). At least one genetic feature determined stimulus response for 10/17 stimuli and at least two features for 6/17 stimuli (Student’s t-tests, FDR = 10%), indicating that CLL viability is controlled both by cell-intrinsic mutations and the cells’ microenvironment. The most prominent factors were IGHV status and trisomy 12. Responses to microenvironmental stimuli were largely independent of receptor expression (Supp. Fig. 7).

To address the possible interplay of multiple genetic factors, we applied linear regression with lasso regularisation to derive for each stimulus a multivariate predictor composed of genetic, IGHV and DNA methylation covariates (Fig. 3B). This revealed at least one genetic predictor for five of the 17 stimuli. Trisomy 12 and IGHV status were again the most common features. Stimulus responses stratified by mutations, along with regression model fits can be explored online (www.dietrichlab.de/CLL_Microenvironment/).

### Trisomy 12 is a key modulator of responses to stimuli

Our survey of genetic determinants of stimuli response highlighted trisomy 12 as a modulator of responses to IL4, TGFβ, soluble CD40L + IL4 and TLR stimuli.

TLR response has previously been shown to depend on the IGHV mutation status^16^; we identified trisomy 12 as a second major determinant of response to TLR stimulation. IGHV mutated CLL samples with trisomy 12 showed a strongly increased viability after TLR stimulation with resiquimod compared to those without trisomy 12. IGHV unmutated CLL cells showed a strong increase in viability upon TLR stimulation regardless of trisomy 12 status (Fig. 4A). In line with our previous findings, we found a trend towards higher toxicity of BCR inhibition in trisomy 12 patients, indicating a dependency of the trisomy 12 effect on the BCR pathway (Fig. 4B)^20^. Further, we observed that the pro-survival effect of TLR stimulation, which is increased in IGHV-U and trisomy 12 CLL samples, could be reversed by BCR inhibition (Supp. Fig. 8)

To investigate the incompletely understood role of trisomy 12 in CLL, we investigated how it might modulate responses to cell-extrinsic signals. We began by investigating the transcriptomic signature of trisomy 12 and performed a gene set enrichment analysis of trisomy 12 vs. non-trisomy 12 samples. Using the hallmark gene sets we found multiple microenvironmental signalling cascades to be upregulated including TNFα and Interferon response pathways (Fig. 4C). Since important microenvironmental pathways are missing in the hallmark gene sets, we repeated this analysis for selected KEGG gene sets relating to microenvironmental signalling and found the BCR and chemokine signalling pathways to be upregulated (Fig. 4D). Overall, trisomy 12 CLL samples exhibit a higher activity of microenvironmental signalling pathways including BCR signalling.

To investigate potential downstream effectors in trisomy 12 CLL, we visualised transcription factor (TF) activity profiles based on chromatin accessibility data derived from ATAC sequencing. We used the diffTF^32^ software to identify TFs with differential binding site accessibility between trisomy 12 and non-trisomy 12 CLL, using ATAC sequencing data for two non-trisomy 12 and two trisomy 12 CLL PBMC samples. The binding sites of 92 TFs respectively were more accessible (p<0.05) in the trisomy 12 samples (Supp. Fig. 9), reflecting a specific signalling signature in trisomy 12 CLL. The top hit was the hematopoietic regulator Spi-B, followed by PU.1, which share similar binding motifs and exhibit functional redundancy^37^. We validated this finding in a second independent dataset taken from Rendeiro et al.^31^, which included 43 WT and nine trisomy 12 samples. The binding sites of nine TFs were more accessible (p<0.05) in the trisomy 12 samples (Fig. 4E), and Spi-B and PU.1 were again the top hits. These TFs are known to be key regulators of healthy B-cell function^38, 39^, controlling B-cell responses to environmental cues including CD40L, TLR ligands and IL4^40^.

We hypothesised that Spi-B might modulate proliferation by coordinating transcriptional response to microenvironmental signals. To identify Spi-B target genes, we reanalysed a ChIPseq dataset^34^, quantifying binding in lymphoma cell lines and tested functional enrichment of immune signalling pathways amongst Spi-B targets. Using selected KEGG and Reactome genesets, we found that TLR, BCR and TGFβ signalling genes were enriched (p<0.01) amongst Spi-B targets (Fig. 4F).

We next investigated whether Spi-B activity and the trisomy 12 signalling signature might be targetable. Amongst the panel of drugs included in this screen, we noted that the bromodomain inhibitor IBET-762 demonstrated higher efficacy in trisomy 12 CLL cells (Fig. 4G). We generated an additional ATACseq dataset consisting of four CLL PBMC samples each treated with IBET-762 and DMSO as control. Using diffTF, we quantified TF binding site accessibility and visualised inferred TF activity for trisomy 12 signature TFs. All nine TFs that showed higher accessibility in trisomy 12 exhibited decreased accessibility upon treatment with IBET-762 (Fig. 4H).

Taken together, we highlight trisomy 12 as a modulator of response to microenvironmental signals, link these effects to differentially active BCR signalling and suggest bromodomain inhibition as a possible therapeutic target in trisomy 12 patients.

### Mapping drug-microenvironment interactions reveals drug resistance and sensitivity pathways

Guided by the observation of reduced treatment efficacy in protective niches^13^, we examined the effects of the stimuli on drug response.

We fitted a linear model (Eqn 1) to quantify how each stimulus modulates drug efficacy beyond its individual effects, such as the impact of the stimuli on baseline viability and spontaneous apoptosis.*β_int_*, termed the interaction factor, quantifies how the combined treatment effect differs from the sum of the individual treatment effects.

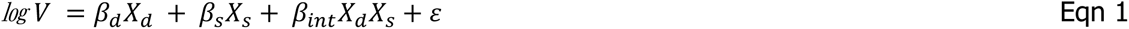

*V* represents the viability with a given treatment, *β_d_*, *β_s_* and *β_int_* are coefficients for the drug, stimulus, and combinatorial terms respectively, *X_d_* and *X_s_* are indicator variables (0 or 1) for the presence or absence of a drug/stimulus. ε is the vector of model residuals.

45 out of 204 combinations had a *β_int_* with p<0.05. We classified these into four categories, based on the sign of *β_int_* and whether the combination effect was antagonistic or synergistic (Fig. 5A-C).

Positive antagonistic interactions were the most common category, in which stimuli reversed drug action and increased viability. These interactions could lead to treatment resistance in vivo. They comprised known resistance mechanisms including the inactivation of ibrutinib by IL4 signalling (Fig. 5D)^18^, and previously unknown resistance pathways, notably the effect of interferon-γ (IFNγ) signalling on ibrutinib response (Fig. 5E).

6/45 interactions were categorised as negative antagonistic, in which drug action reversed the pro-survival effect of microenvironmental stimulation. These interactions point to strategies to overcome treatment resistance. For instance, Pan-JAK inhibition reduced the increase in viability from sCD40L + IL4 stimulation (Fig. 5F).

We observed a single positive synergistic case, whereby simultaneous treatment with IFNγ and ralimetinib, a p38 MAPK inhibitor, induced a synergistic increase in viability which was not observed with either single treatment (Fig. 5G), pointing towards an inhibitory effect of p38 activity on IFNγ signalling.

16 combinations showed negative synergistic interactions. For example, TLR9 agonist CpG ODN increased the efficacy of BCR inhibition (Fig. 5H). Interestingly, in samples where TLR stimulation increased viability, the addition of BCR inhibitors (ibrutinib and idelalisib) suppressed this effect, indicating that the increase of viability upon TLR stimulation is dependent on BCR activity. The efficacy of luminespib, a HSP90 inhibitor, increased with various stimuli, including soluble anti-IgM (Fig. 5I).

Altogether, these results highlight the influence of the microenvironment on drug efficacy and underline the value of microenvironmental stimulation in ex vivo studies of drug efficacy.

### Genetic features affect drug-microenvironment interactions

Next, to investigate to what extent genetic driver mutations modulated the interactions between stimuli and drugs, we fit the linear model in Eqn. (1) in a sample - specific manner.

This resulted in a sample-specific *β_int_* interaction coefficient for each drug - stimulus combination. We looked for associations between the size of *β_int_* and genetic features using multivariate regression with L1 (lasso) regularisation, with gene mutations and IGHV status as predictors. We generated a predictor profile for each drug - stimulus combination, considering predictors that were selected in >90% of bootstrapped model fits. In total, we found genetic modulators for 60/204 interactions (Fig 6A, Supp Fig. 10). Trisomy 12 and IGHV status modulated the largest number of interactions.

**Figure 6.**
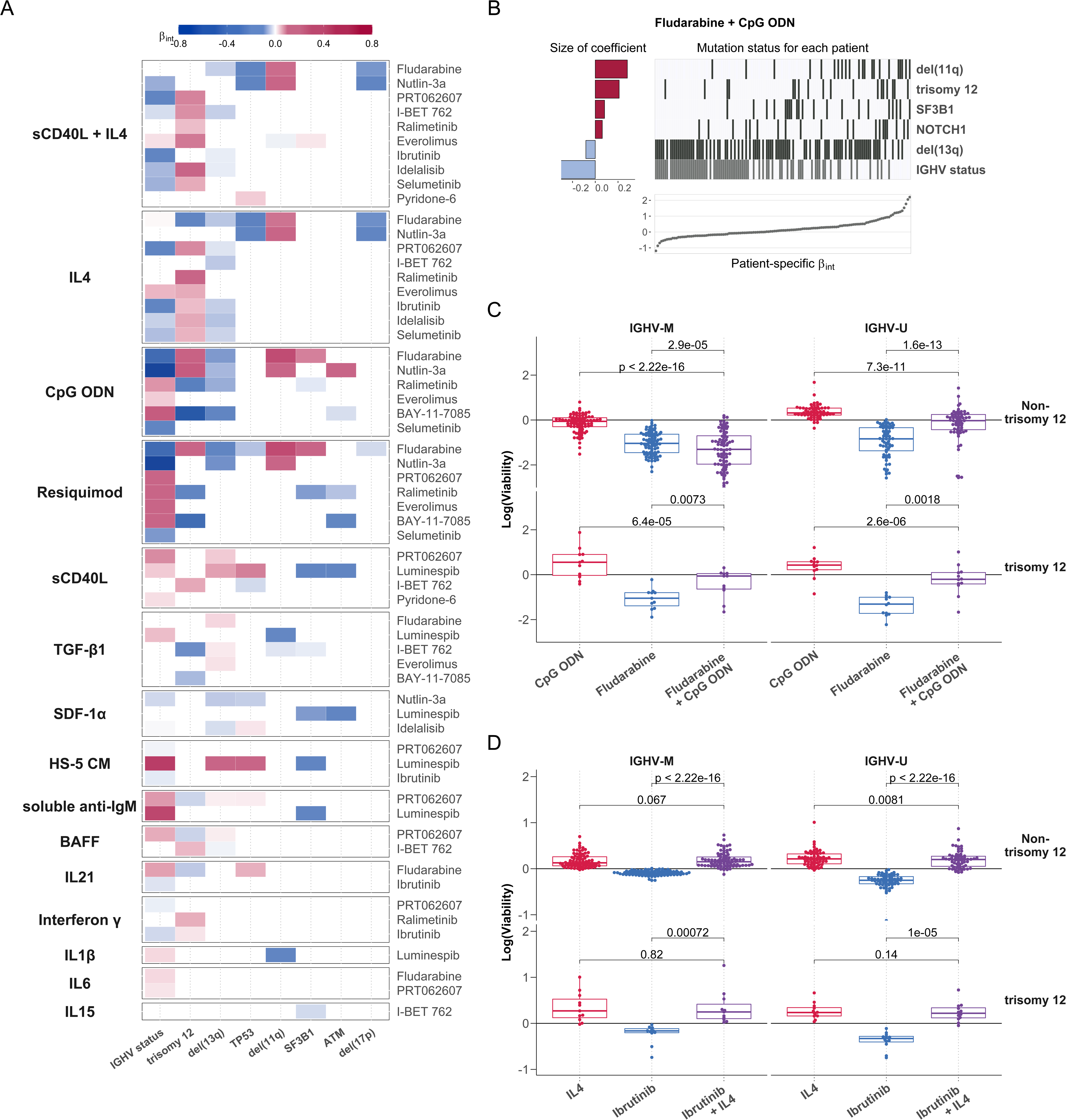
Integrating the effects of genetic features and stimuli on drug response. (A) Heatmap depicting the eight most commonly selected genetic predictors of drug-stimulus interactions (each row represents the coefficients of a single multivariate model, as in (B)). Stimuli are shown on left, and corresponding drugs on right. Coloured fields indicate that βint for given drug and stimulus is modulated by corresponding genetic feature. Positive coefficients are shown in red, indicating a more positive *β_int_* if the feature is present. Drug-stimulus combinations with no genetic predictors of *β_int_* amongst shown genetic factors are omitted for clarity. (B) Predictor profile depicting genetic features that modulate the interaction between fludarabine and CpG ODN. The horizontal bars on left show the size of fitted coefficients assigned to genetic features. The matrix in the centre indicates patient mutation status for the selected genetic features aligned with the scatter plot indicating the size of βint for each patient. Grey lines indicate the presence of genetic feature/IGHV mutated. (C-D) Beeswarm boxplots of log(viability) values, for fludarabine + CpG ODN (C) and ibrutinib + IL4 (D) single and combinatorial treatments, faceted by IGHV status and trisomy 12 status. P-values from paired Student’s t-tests.

For example, we found the value of *β_int_* for fludarabine and CpG ODN to be modulated by six genetic factors; most strongly by IGHV status, del(11q), and trisomy 12 (Fig. 6B).

In IGHV-M non-trisomy 12 samples, fludarabine efficacy increased in the presence of TLR stimulation, whilst in the other subgroups TLR stimulation induced resistance to fludarabine (Fig. 6C).

Our approach also identified interactions dependent on single treatment effects. IL4 induced complete resistance to ibrutinib regardless of higher ibrutinib efficacy in trisomy 12 and IGHV- U samples. This highlights the breadth of IL4-induced resistance to ibrutinib in CLL across genetic backgrounds (Fig. 6D, Supp. Fig. 11).

Overall, these findings illustrate how the presence of known genetic alterations determines how drugs and external stimuli interact with each other.

### The key resistance pathways IL4 and TLR show increased activity in CLL-infiltrated lymph nodes

Microenvironmental signalling within lymph nodes has been implicated in treatment resistance^9, 41–43^, though a clear understanding of the mechanisms involved remains missing. Since IL4 and TLR signalling were the most prominent modulators of drug response in our study, we assessed their activity within the lymph node niche. We stained paraffin embedded sections of 100 CLL-infiltrated and 100 non-neoplastic lymph nodes for pSTAT6, an essential downstream target of IL4 (Supp. Fig. 12), and pIRAK4, a downstream target of TLR7/8/9.

The CLL-infiltrated lymph nodes showed higher levels of pSTAT6 (Fig. 7A) and pIRAK4 (Fig. 7B). Examples are shown in Fig. 7C-F.

**Figure 7.**
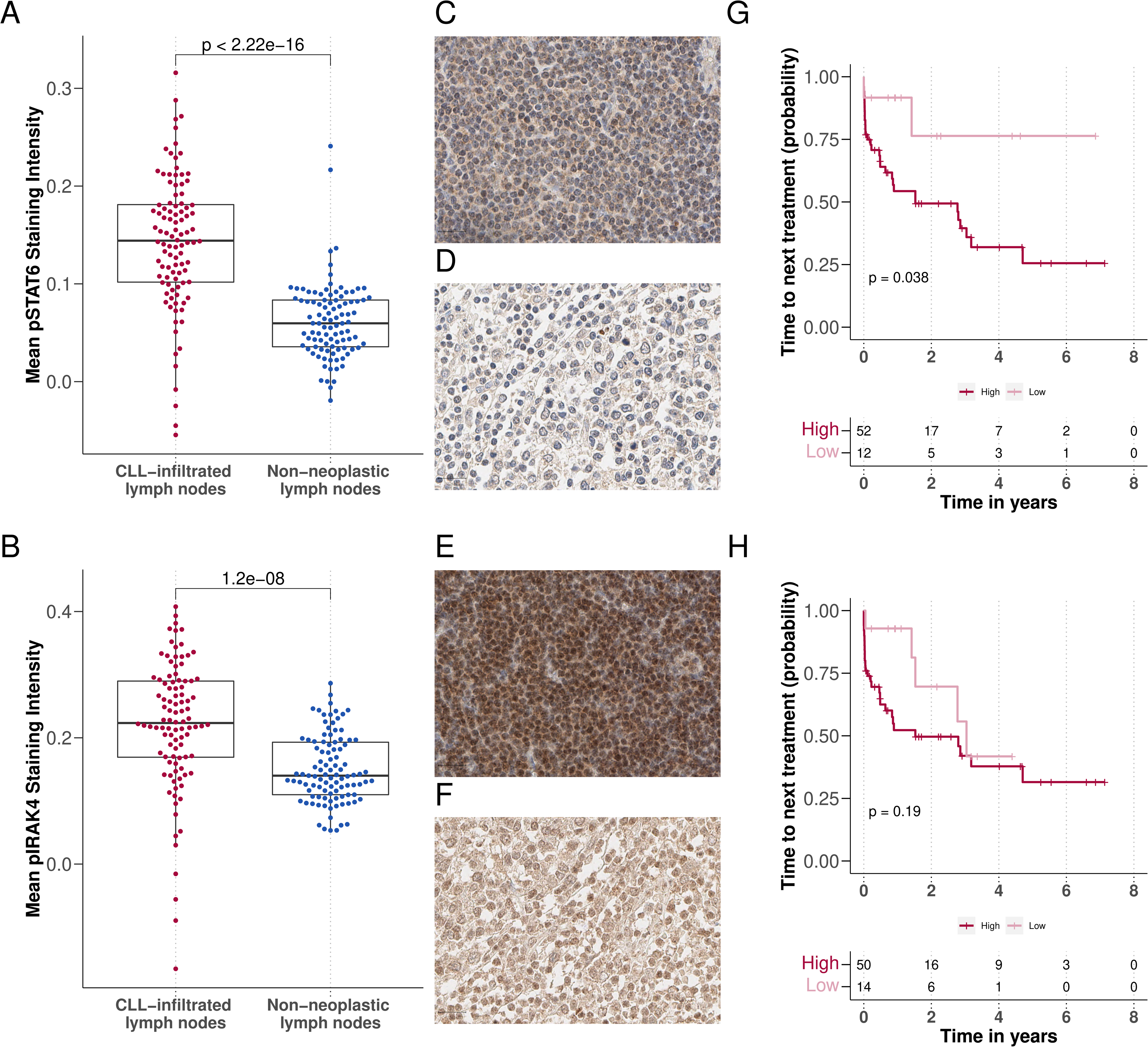
IL4 and TLR signalling are upregulated in CLL-infiltrated lymph nodes. (A-B) Mean pSTAT6 (A) and pIRAK4 (B) staining intensity in CLL-infiltrated and non-neoplastic lymph node biopsies after background subtraction (y axis), p-values from Student’s t-test. Each dot represents the mean of all cells in TMA cores per patient sample. (C-F) Example images of IHC sections. (C + D) show pSTAT6 levels in (C) CLL-infiltrated and (D) non-neoplastic samples. (E + F) show pIRAK4 levels in (E) CLL-infiltrated and (F) non-neoplastic samples. (G + H) Kaplan-Meier plots for time to next treatment stratified by levels (high / low) of pSTAT6 (G) and pIRAK4 (H).

To investigate the influence of microenvironmental activity in lymph nodes on CLL disease progression, we correlated staining intensity to time to next treatment. High activity of pSTAT- 6 correlated with shorter time to next treatment (Fig. 7G), the same trend could be observed with higher pIRAK4 (Fig. 7H). Higher IL4 activity within the lymph node appeared to relate to shorter time to next treatment, and may provide further evidence to the hypothesis that microenvironmental activity promotes treatment resistance within the lymph node, eventually leading to relapse.

## Discussion

Our work maps the effects of microenvironmental stimuli in the presence of drugs and links these to underlying molecular properties across 192 primary CLL samples. We employ all combinations of 17 stimuli with 12 drugs as a reductionist model of microenvironmental signalling. We account for the confounding effects of spontaneous apoptosis ex vivo and dissect the effect of individual microenvironmental stimuli on baseline viability and drug toxicity. The results may serve as building blocks for a more holistic understanding of the interactions of tumour genetics, microenvironment, and drug response in complex in vivo situations.

We discover that CLL subgroups can be extracted from microenvironmental response phenotypes and that this classification is linked to distinct molecular profiles and clinical outcomes. IGHV-M samples with weaker responses to microenvironmental stimuli showed faster disease progression, highlighting the important role microenvironmental signalling may play in CLL pathophysiology.

The sensitivity afforded by our approach enabled us to identify drug-stimulus interactions, meaning instances where the effect of a drug on the tumour cells is modulated by a microenvironmental stimulus. Our assay recapitulated known drug-resistance phenomena, such as the impact of IL4 stimulation on ibrutinib and idelalisib^18^ and identified new ones, including IFNγ-induced resistance to BCR inhibition. Systematic mapping of drug-stimulus interactions, such as generated in this study, can be used to inform biology-based combinatorial treatments across different entities.

We demonstrated the breadth of IL4-induced resistance against kinase inhibitors and chemotherapeutics across a range of genetic backgrounds. IL4 induced complete resistance to BCR inhibitors across genetic subgroups, despite greater BCR inhibitor efficacy in IGHV-U CLL. Further, we observed increased IL4 activity within CLL-infiltrated lymph nodes and the association of higher activity with faster disease progression. Our work highlights the significance of IL4 within the lymph node niche, especially since proliferation^9^ and resistance to BCR inhibition^13^ is linked to lymph nodes and our recent work suggests T follicular helper cells as a source of IL4^44^. Targeting pathways downstream of IL4, via JAK inhibition, has been reported to improve BCR response in non-responding CLL patients^45^.

Underneath cell-extrinsic factors that control the apoptotic and proliferative activity of tumour cells in response to drugs there lies the network of cell-intrinsic molecular components. We used multivariate modelling to capture the impact of molecular features on responses to stimuli and drugs. For instance, the response to TLR stimulation depended on IGHV status, trisomy 12, del11q, del13q and ATM, reflecting the multiple layers of biology involved. Most strikingly, we observed that TLR stimulation increased fludarabine toxicity in the subgroup of IGHV-M non-trisomy 12 patient samples, while it led to fludarabine resistance in all other backgrounds. This finding combined with our observation that TLR signalling is highly active in CLL-infiltrated lymph nodes might help explain the heterogeneous effects of chemotherapy in CLL, in particular that fludarabine therapy can achieve lasting remission in IGHV-M but not in IGHV- U patients.

Beyond TLR, trisomy 12 modulated a broad range of stimulus responses and drug-stimulus interactions, comparable to the impact of IGHV status. Molecularly, we link the observed trisomy 12 phenotype to upregulated microenvironmental signalling and higher activity of Spi- B, an essential regulator of environmental sensing in B-cells^40^. The TF signature associated with trisomy 12 can be reversed by bromodomain inhibition and trisomy 12 samples exhibit increased sensitivity to bromodomain inhibition, indicating this as potential therapeutic strategy in trisomy 12 CLL.

We present a data resource for the study of drug response in the context of cell-intrinsic and cell-extrinsic modulators. The data may inform targeted mechanistic investigations^30^ and direct efforts for combination therapies. The entire dataset, including the reproducible analysis, can be downloaded from the online repository (github.com/Huber-group-EMBL/CLLCytokineScreen2021), or explored interactively (www.dietrichlab.de/CLL_Microenvironment/).

## Acknowledgments

We thank the Department of Medicine V, University of Heidelberg, Molecular Medicine Partnership Unit (MMPU), EMBL Heidelberg and DKFZ/Heidelberg Center for Personalized Oncology (DKFZ/HIPO) for technical support and funding. H.A.R.G. was supported by a Joachim-Herz Add-on Fellowship for Interdisciplinary Science. P.M.B was supported by a doctoral scholarship of the Franziska-Kolb foundation. S.D. was supported by the Else Kröner Fresenius Foundation, the Federal Ministry of Education and Research (SYMPATHY) and the German Research Foundation (SFB873). J.L., S.D. and W.H. were supported by the German Federal Ministry of Education and Research (CompLS project MOFA under grant agreement number 031L0171A). T.R. was supported by a physician scientist fellowship of the Medical Faculty of University Heidelberg and the German Federal Ministry of Education and Research (CompLS project SIMONA under grant agreement number 031L0263A). For experimental technical support and expertise, we thank Mareike Knoll, Angela Lenze, Brian Lai, and Vladimir Benes, and Nayara Trevisan Doimo de Azevedohe of the EMBL Genomics Core Facility. We thank EMBL’s IT Services for providing data management and computing infrastructure and Simone Bell for organisational support. We thank Mike Smith and Frederik Ziebell for advice on statistics and analysis, Martina Seiffert for the HS-5 stromal cell line, and Michael Boutros, Moritz Gerstung, Nassos Typas, and Simon Anders for their comments on the manuscript and their valued advice. P.M.B is a MD candidate at the University of Heidelberg. This work is submitted in partial fulfilment of the requirement for the MD. H.A.R.G is a PhD candidate for the joint PhD degree between EMBL and Heidelberg University, Faculty of Biosciences. This work is submitted in partial fulfilment of the requirement for the PhD.

## Author contributions

S.D. and P.M.B. designed the experiments. S.D., W.H., H.A.R.G., and P.M.B. conceptualised the data analysis. W.H. and S.D. supervised the study. W.H., T.Z. and S.D. conceptualised the multi-omics approach to analyse CLL drug response. P.M.B. performed the drug-stimulus profiling experiments with support from C.K., S.S., and L.W.. T.Z., C.M.T., and P.D. provided guidance on experimental protocols. H.A.R.G., P.M.B, J.H., J.L., S.D. performed data processing and curation. H.A.R.G., and P.M.B. analysed and interpreted the data, with support from W.H., S.D., J.L, S.H., and T.R., and produced the figures with input from W.H. and S.D.. H.A.R.G. performed statistical inference and regression modelling, P.M.B. performed exploratory analysis and integration with clinical data. T.B. and S.H. performed the shRNA knockdown experiments, J.B.Z. and I.B. provided the ATACseq dataset used in Figure 4 and quantified differential transcription factor activity, M.K., K.K., and C.Z. generated the immunohistochemistry data, and P.M.B. analysed these data. H.A.R.G. designed and wrote the Shiny app and the software documentation. P.M.B. and H.A.R.G. wrote the manuscript, with input from S.D. and W.H.. All authors reviewed and edited the manuscript.

## References

1. Fabbri, G. & Dalla-Favera, R. The molecular pathogenesis of chronic lymphocytic leukaemia. Nat. Rev. Cancer 16, 145–162 (2016).

2. Puente, X. S., Jares, P. & Campo, E. Chronic lymphocytic leukemia and mantle cell lymphoma: crossroads of genetic and microenvironment interactions. Blood 131, 2283– 2296 (2018).

3. Burger, J. A. et al. Ibrutinib as Initial Therapy for Patients with Chronic Lymphocytic Leukemia. N. Engl. J. Med. 373, 2425–2437 (2015).

4. Fischer, K. et al. Venetoclax and Obinutuzumab in Patients with CLL and Coexisting Conditions. N. Engl. J. Med. 380, 2225–2236 (2019).

5. Gaidano, G. & Rossi, D. The mutational landscape of chronic lymphocytic leukemia and its impact on prognosis and treatment. Hematology Am. Soc. Hematol. Educ. Program 2017, 329–337 (2017).

6. Döhner, H. et al. Genomic aberrations and survival in chronic lymphocytic leukemia. N. Engl. J. Med. 343, 1910–1916 (2000).

7. Strefford, J. C. The genomic landscape of chronic lymphocytic leukaemia: biological and clinical implications. Br. J. Haematol. 169, 14–31 (2015).

8. Burger, J. A. et al. Long-term efficacy and safety of first-line ibrutinib treatment for patients with CLL/SLL: 5 years of follow-up from the phase 3 RESONATE-2 study. Leukemia 34, 787–798 (2020).

9. Herndon, T. M. et al. Direct in vivo evidence for increased proliferation of CLL cells in lymph nodes compared to bone marrow and peripheral blood. Leukemia 31, 1340–1347 (2017).

10. Collins, R. J. et al. Spontaneous programmed death (apoptosis) of B-chronic lymphocytic leukaemia cells following their culture in vitro. Br. J. Haematol. 71, 343–350 (1989).

11. Oppermann, S. et al. High-content screening identifies kinase inhibitors that overcome venetoclax resistance in activated CLL cells. Blood 128, 934–947 (2016).

12. Ten Hacken, E. & Burger, J. A. Microenvironment interactions and B-cell receptor signaling in Chronic Lymphocytic Leukemia: Implications for disease pathogenesis and treatment. Biochim. Biophys. Acta 1863, 401–413 (2016).

13. Ahn, I. E. et al. Depth and durability of response to ibrutinib in CLL: 5-year follow-up of a phase 2 study. Blood 131, 2357–2366 (2018).

14. Chen, Z. et al. LINKING MICROENVIRONMENTAL SIGNALS TO METABOLIC SWITCHES AND IBRUTINIB RESPONSE IN CHRONIC LYMPHOCYTIC LEUKEMIA: PS1125. HemaSphere 3, 509–510 (2019).

15. Jayappa, K. D. et al. Microenvironmental agonists generate phenotypic resistance to combined ibrutinib plus venetoclax in CLL and MCL. Blood Adv 1, 933–946 (2017).

16. Chatzouli, M. et al. Heterogeneous functional effects of concomitant B cell receptor and TLR stimulation in chronic lymphocytic leukemia with mutated versus unmutated Ig genes. J. Immunol. 192, 4518–4524 (2014).

17. Dietrich, S. et al. Leflunomide induces apoptosis in fludarabine-resistant and clinically refractory CLL cells. Clin. Cancer Res. 18, 417–431 (2012).

18. Aguilar-Hernandez, M. M. et al. IL-4 enhances expression and function of surface IgM in CLL cells. Blood 127, 3015–3025 (2016).

19. Carey, A. et al. Identification of Interleukin-1 by Functional Screening as a Key Mediator of Cellular Expansion and Disease Progression in Acute Myeloid Leukemia. Cell Rep. 18, 3204–3218 (2017).

20. Dietrich, S. et al. Drug-perturbation-based stratification of blood cancer. J. Clin. Invest. 128, 427–445 (2018).

21. Bankhead, P. et al. QuPath: Open source software for digital pathology image analysis. Sci. Rep. 7, 1–7 (2017).

22. Love, M. I., Huber, W. & Anders, S. Moderated estimation of fold change and dispersion for RNA-seq data with DESeq2. Genome Biol. 15, 550 (2014).

23. Therneau, T. M. & Grambsch, P. M. Modeling Survival Data: Extending the Cox Model. (Springer Science & Business Media, 2000).

24. Friedman, J., Hastie, T. & Tibshirani, R. Regularization Paths for Generalized Linear Models via Coordinate Descent. J. Stat. Softw. 33, 1–22 (2010).

25. Wilkerson, M. D. & Hayes, D. N. ConsensusClusterPlus: a class discovery tool with confidence assessments and item tracking. Bioinformatics 26, 1572–1573 (2010).

26. Yu, G., Wang, L.-G., Han, Y. & He, Q.-Y. clusterProfiler: an R Package for Comparing Biological Themes Among Gene Clusters. OMICS: A Journal of Integrative Biology vol. 16 284–287 (2012).

27. Yu, G., Wang, L.-G. & He, Q.-Y. ChIPseeker: an R/Bioconductor package for ChIP peak annotation, comparison and visualization. Bioinformatics 31, 2382–2383 (2015).

28. Akalin, A., Franke, V., Vlahoviček, K., Mason, C. E. & Schübeler, D. Genomation: a toolkit to summarize, annotate and visualize genomic intervals. Bioinformatics 31, 1127–1129 (2015).

29. Malgorzata Oles, Sascha Dietrich, Junyan Lu, Britta Velten, Andreas Mock, Vladislav Kim, Wolfgang Huber. BloodCancerMultiOmics2017. (Bioconductor, 2018). doi:10.18129/B9.BIOC.BLOODCANCERMULTIOMICS2017.

30. Lu, J. et al. Multi-omics reveals clinically relevant proliferative drive associated with mTOR-MYC-OXPHOS activity in chronic lymphocytic leukemia. Nat Cancer (2021) doi:10.1038/s43018-021-00216-6.

31. Rendeiro, A. F. et al. Chromatin accessibility maps of chronic lymphocytic leukaemia identify subtype-specific epigenome signatures and transcription regulatory networks. Nat. Commun. 7, 11938 (2016).

32. Berest, I. et al. Quantification of Differential Transcription Factor Activity and Multiomics-Based Classification into Activators and Repressors: diffTF. Cell Rep. 29, 3147–3159.e12 (2019).

33. Kulakovskiy, I. V. et al. HOCOMOCO: expansion and enhancement of the collection of transcription factor binding sites models. Nucleic Acids Res. 44, D116–25 (2016).

34. Care, M. A. et al. SPIB and BATF provide alternate determinants of IRF4 occupancy in diffuse large B-cell lymphoma linked to disease heterogeneity. Nucleic Acids Res. 42, 7591–7610 (2014).

35. Edgar, R., Domrachev, M. & Lash, A. E. Gene Expression Omnibus: NCBI gene expression and hybridization array data repository. Nucleic Acids Res. 30, 207–210 (2002).

36. Baumann, T. et al. Lymphocyte doubling time in chronic lymphocytic leukemia modern era: a real-life study in 848 unselected patients. Leukemia (2021) doi:10.1038/s41375-021-01149-w.

37. Garrett-Sinha, L. A., Dahl, R., Rao, S., Barton, K. P. & Simon, M. C. PU.1 exhibits partial functional redundancy with Spi-B, but not with Ets-1 or Elf-1. Blood 97, 2908–2912 (2001).

38. Turkistany, S. A. & DeKoter, R. P. The Transcription Factor PU.1 is a Critical Regulator of Cellular Communication in the Immune System. Arch. Immunol. Ther. Exp. 59, 431– 440 (2011).

39. Ray-Gallet, D., Mao, C., Tavitian, A. & Moreau-Gachelin, F. DNA binding specificities of Spi-1/PU.1 and Spi-B transcription factors and identification of a Spi-1/Spi-B binding site in the c-fes/c-fps promoter. Oncogene 11, 303–313 (1995).

40. Willis, S. N. et al. Environmental sensing by mature B cells is controlled by the transcription factors PU.1 and SpiB. Nat. Commun. 8, 1426 (2017).

41. Hayden, R. E., Pratt, G., Roberts, C., Drayson, M. T. & Bunce, C. M. Treatment of chronic lymphocytic leukemia requires targeting of the protective lymph node environment with novel therapeutic approaches. Leuk. Lymphoma 53, 537–549 (2012).

42. Mittal, A. K. et al. Chronic lymphocytic leukemia cells in a lymph node microenvironment depict molecular signature associated with an aggressive disease. Mol. Med. 20, 290– 301 (2014).

43. Yan, X.-J. et al. Identification of outcome-correlated cytokine clusters in chronic lymphocytic leukemia. Blood 118, 5201–5210 (2011).

44. Roider, T. et al. Dissecting intratumour heterogeneity of nodal B-cell lymphomas at the transcriptional, genetic and drug-response levels. Nature Cell Biology vol. 22 896–906 (2020).

45. Spaner, D. E., McCaw, L., Wang, G., Tsui, H. & Shi, Y. Persistent janus kinase-signaling in chronic lymphocytic leukemia patients on ibrutinib: Results of a phase I trial. Cancer Med. 8, 1540–1550 (2019).

