## Supplemental Material for "Combinatorial drug-microenvironment interaction mapping reveals cell-extrinsic drug resistance mechanisms and clinically relevant patient subgroups in CLL"

9th November 2021

### Supplementary Methods

#### Drug-stimulation profiling of samples treated with Ibrutinib, IL4 and AS1517499

The experiment in Supp. Fig. 11, was performed as detailed in the main methods (Sample preparation and drug-stimulation profiling), but with the following adjustments. 16 independent CLL PBMC samples were used and luminescence was read out using a Perkin Elmer EnSight.

#### ATAC sequencing

Peripheral blood was taken from 4 CLL patients and separated by Ficoll gradient (GE Healthcare), mononuclear cells were cryopreserved on liquid nitrogen. Samples were later thawed from frozen as previously described[1] and MACS sorted for CD19 positive cells (Miltenyi autoMACS®). The cells were resuspended in RPMI (GIBCO, Cat.No. 21875–034), with the addition of 2mM glutamine (GIBCO, Cat.No. 25030–24), 1% Pen/Strep (GIBCO, Cat.No. 15140–122) and 10% pooled, heat–inactivated and sterile filtered human type AB male off the clot serum (PAN Biotech, Cat.No. P40–2701, Lot.No:P–020317). 5ml of cell suspension was cultured in 6–well plates (Greiner Bio–One Cat.No. 657160). After thawing, cells were incubated at 37°C and 5% $CO_2$ for 6 hours in 0.2% DMSO. The final cell concentration was 2x10ˆ6 cells/ml. Cell viability and purity was assessed using FACS. All samples had a viability over 90% and over 95% of CD19+/CD5+/CD3– cells.

#### ATAC sequencing library generation

ATACseq libraries were generated as described previously[2]. Cell preparation and transposition was performed according to the protocol, starting with 5x10ˆ4 cells per sample. Purified DNA was stored at −20°C until library preparation was performed. To generate multiplexed libraries, the transposed DNA was initially amplified for 5x PCR cycles using 2.5 μL each of 25 μM PCR Primer 1 and 2.5 μL of 25 μM Barcoded PCR Primer 2 (included in the Nextera index kit, Illumina, San Diego, CA, USA), 25 μL of NEBNext High-Fidelity 2x PCR Master Mix (New England Biolabs, Boston, Massachusetts) in a total volume of 50 μL. 5 μL of the amplified DNA was used to determine the appropriate number of additional PCR cycles using qPCR. Additional number of cycles was calculated through the plotting of the linear Rn versus cycle, and corresponds to one-third of the maximum fluorescent intensity. Finally, amplification was performed on the remaining 45 μL of the PCR reaction using the optimal number of cycles determined for each library by qPCR (max. 13 cycles in total). The amplified fragments were purified with two rounds of SPRI bead clean-up (1.4x). The size distribution of the libraries was assessed on Bioanalyzer with a DNA High Sensitivity kit (Agilent Technologies, Santa Clara, CA), concentration was measured with Qubit® DNA High Sensitivity kit in Qubit® 2.0 Flurometer (Life Technologies, Carlsbad, CA). Sequencing was performed on NextSeq 500 (Illumina, San Diego, CA, USA) using 75bp paired-end sequencing, generating approx. 450 million paired-reads per run, with an average of 55 million reads per sample.

#### ATAC sequencing analysis of transcription factor activity in trisomy 12 CLL

Raw ATACseq data generated from our CLL samples were processed as described in Berest et al. 2019[3], with the only exception that we did not use CG bias correction step. We then used analytical mode of diffTF with HOCOMOCO v10 database[4] using the following parameters: minOverlap = 1; design formula = "~ Patient + trisomy 12 status".

| Drug | Main targets | Target category | Drug Group | Pathway | Distributor | Catalogue number | Conc. 1 | Conc. 2 |
|---|---|---|---|---|---|---|---|---|
| Ibrutinib | BTK | BCR | kinase inhibitor | B-cell receptor | Selleck Chemicals | S2680 | 500nM | 50nM |
| Idelalisib | PI3K delta | BCR | kinase inhibitor | B-cell receptor | Selleck Chemicals | S2226 | 500nM | 50nM |
| Fludarabine | Purine analogue | DDR | chemotherapeutic agent | Other | Selleck Chemicals | S1491 | 2000nM | 200nM |
| Nutlin-3a | MDM2 | DDR | other | Other | Selleck Chemicals | S8059 | 10000nM | 1000nM |
| Selumetinib | MEK1/2 | MAPK | kinase inhibitor | Other | Selleck Chemicals | S1008 | 1000nM | 100nM |
| BAY-11-7085 | NFkB | NFkB | other | Other | Selleck Chemicals | S7352 | 2000nM | 200nM |
| Everolimus | mTOR | mTOR | other | Other | Selleck Chemicals | S1120 | 500nM | 50nM |
| PRT062607 | SYK | BCR | kinase inhibitor | B-cell receptor | Selleck Chemicals | S8032 | 500nM | 50nM |
| Pyridone-6 | JAK1/2/3 | JAK/STAT | kinase inhibitor | Other | MedChemExpress | 457021-03-7 | 500nM | 50nM |
| Ralimetinib | p38 MAPK | MAPK | kinase inhibitor | Other | Selleck Chemicals | S1494 | 1500nM | 150nM |
| Luminespib | HSP90 | HSP90 | other | Other | Selleck Chemicals | S1069 | 200nM | 20nM |
| I-BET 762 | BRD2/3/4 | Epigenome | other | Other | Selleck Chemicals | S7189 | 1000nM | 100nM |

**Supplementary Table 1. Drugs contained in the study.** List of drugs, along with their main target, supplier and concentration. All drugs were dissolved in Dimethyl Sulfoxide (DMSO) and stored at –20°C.

| Stimulus | Name | Supplier | Concentration | Catalogue Number | Lot Number | Pathway |
|---|---|---|---|---|---|---|
| IL4 human recombinant animal component free | IL4 | Sigma-Aldrich | 10 ng/ml | SRP3093 | 0712AFC14 | JAK/STAT |
| IL10 human Animal component free | IL10 | Sigma-Aldrich | 10 ng/ml | SRP3312 | 1012AFC21 | JAK/STAT |
| IL2 human recombinant animal component free | IL2 | Sigma-Aldrich | 10 ng/ml | SRP3085 | 0416AFC12 | JAK/STAT |
| R-848 | Resiquimod | Enzo Life Siences | 1000 ng/ml | ALX-420-038-M025 | 10211615 | TLR 7/8 |
| Human IL-21 | IL21 | Peprotech | 10 ng/ml | 200-21 | 414226 | JAK/STAT |
| Human BAFF | BAFF | Peprotech | 250 ng/ml | 310-13 | 0706CY194 | NFkB |
| Human IL-1 beta | IL1ß | Peprotech | 10 ng/ml | 200-01 | 0606B95 | NFkB |
| Human sCD40 Ligand | sCD40L | Peprotech | 1000 ng/ml | 310-02 | 1214145 | NFkB |
| Goat F(AB')2 Fragment to human IgM | soluble anti-IgM | MP Biomedicals | 20000 ng/ml | 55055 | 7227 | BCR |
| Human TGFß1 | TGFß1 | Peprotech | 10 ng/ml | 100-21 | 1117209 | MAPK |
| Human IL15 | IL15 | Peprotech | 10 ng/ml | 200-15 | 91624 | JAK/STAT |
| Human IL6 | IL6 | Peprotech | 10 ng/ml | 200-06 | 031316-2 | JAK/STAT |
| ODN 2006 (ODN 7909) | CpG ODN | Invivogen | 1000 ng/ml | tlrl-2006-1 | 3901-09T | TLR 9 |
| Human SDF1 alpha (CXCL12) | SDF-1a | Peprotech | 200 ng/ml | 300-28A | 101492 | JAK/STAT |
| Human Interferon gamma | Interferon<U+03B3> | Peprotech | 5 ng/ml | 300-02 | 121527 | NFkB |
| HS-5 conditioned medium | HS-5 CM | self produced | 20 % | NA | NA | NA |

**Supplementary Table 2. Stimuli contained in the study.** List of microenvironmental stimuli, supplier, concentration and main pathway.

| Patient ID | Sex | Treated before | IGHV status | Methylation Cluster | Del13q | Del11q | Trisomy 12 | Del17p |
|---|---|---|---|---|---|---|---|---|
| Pat_001 | f | 1 | U | LP | 1 | 0 | 0 | 0 |
| Pat_002 | m | 1 | M | IP | 1 | 0 | 0 | 0 |
| Pat_003 | m | 0 | M | HP | 0 | 0 | 1 | 0 |
| Pat_004 | f | 1 | U | LP | 0 | 1 | 0 | 0 |
| Pat_005 | m | 0 | U | LP | 1 | 0 | 0 | 0 |
| Pat_006 | f | 0 | U | LP | 0 | 0 | 0 | 0 |
| Pat_007 | f | 0 | M | HP | 1 | 0 | 0 | 0 |
| Pat_008 | m | 1 | U | LP | 1 | 0 | 0 | 0 |
| Pat_009 | m | 1 | U | LP | 1 | 0 | 0 | 1 |
| Pat_010 | f | 1 | U | LP | 1 | 0 | 0 | 1 |
| Pat_011 | f | 0 | U | NA | 0 | 0 | 1 | 0 |
| Pat_012 | f | 0 | M | HP | 1 | 0 | 0 | 0 |
| Pat_013 | f | 1 | U | IP | 1 | 1 | 0 | 0 |
| Pat_014 | m | 0 | M | HP | 0 | 0 | 0 | 0 |
| Pat_015 | m | 0 | M | HP | 1 | 0 | 0 | 0 |
| Pat_016 | m | 0 | M | HP | 1 | 0 | 0 | 0 |
| Pat_017 | m | 0 | M | NA | 1 | 0 | 0 | 0 |
| Pat_018 | f | 1 | U | LP | 1 | 1 | 0 | 0 |
| Pat_019 | m | 0 | M | HP | 1 | 0 | 0 | 0 |
| Pat_020 | f | 1 | M | IP | 1 | 0 | 0 | 0 |
| Pat_021 | m | 0 | U | LP | 0 | 0 | 0 | 1 |
| Pat_022 | f | 0 | M | IP | 0 | 0 | 1 | 0 |
| Pat_023 | f | 0 | M | HP | 0 | 0 | 0 | 0 |
| Pat_024 | f | 1 | M | HP | 0 | 0 | 0 | 1 |
| Pat_025 | m | 0 | M | IP | 1 | 0 | 0 | 0 |
| Pat_026 | m | 0 | M | HP | 1 | 0 | 0 | 0 |
| Pat_027 | f | 0 | M | HP | 1 | 0 | 0 | 0 |
| Pat_028 | f | 0 | M | IP | 1 | 0 | 0 | 0 |
| Pat_029 | f | 0 | M | HP | 1 | 0 | 0 | 0 |
| Pat_030 | m | 1 | M | HP | 1 | 0 | 0 | 0 |
| Pat_031 | m | 0 | M | HP | 0 | 0 | 0 | 0 |
| Pat_032 | f | 1 | U | LP | 1 | 1 | 0 | 0 |
| Pat_033 | m | 1 | U | IP | 0 | 1 | 0 | 0 |
| Pat_034 | m | 1 | U | LP | 0 | 1 | 0 | 0 |
| Pat_035 | m | 0 | M | HP | 1 | 0 | 0 | 0 |
| Pat_036 | m | 1 | U | LP | 1 | 1 | 0 | 1 |
| Pat_037 | f | 1 | U | IP | 1 | 0 | 0 | 0 |
| Pat_038 | m | 0 | M | IP | 1 | 0 | 0 | 0 |
| Pat_039 | m | 1 | M | HP | 1 | 0 | 0 | 0 |
| Pat_040 | f | 0 | M | HP | 0 | 0 | 0 | 0 |
| Pat_041 | f | 1 | U | LP | 0 | 0 | 1 | 0 |
| Pat_042 | f | 0 | M | IP | 1 | 1 | 0 | 0 |
| Pat_043 | f | 0 | M | HP | 1 | 0 | 0 | 0 |
| Pat_044 | m | 0 | M | HP | 1 | 0 | 0 | 0 |
| Pat_045 | m | 0 | U | LP | 0 | 0 | 0 | 0 |
| Pat_046 | m | 0 | M | IP | 0 | 0 | 1 | 0 |
| Pat_047 | m | 0 | M | HP | 1 | 0 | 0 | 0 |
| Pat_048 | m | 0 | M | HP | 1 | 0 | 0 | 0 |
| Pat_049 | f | 0 | M | IP | 0 | 1 | 0 | 0 |
| Pat_050 | m | 0 | M | HP | 0 | 0 | 1 | 0 |
| Pat_051 | m | 0 | M | NA | 1 | 0 | 0 | 0 |
| Pat_052 | f | 0 | M | IP | 1 | 0 | 0 | 0 |
| Pat_053 | m | 1 | U | LP | 1 | 1 | 0 | 0 |
| Pat_054 | f | 1 | U | LP | 0 | 1 | 0 | 0 |
| Pat_055 | m | 1 | U | LP | 0 | 0 | 0 | 0 |
| Pat_056 | f | 0 | U | LP | NA | NA | NA | NA |
| Pat_057 | f | 0 | M | HP | 1 | 0 | 1 | 0 |
| Pat_058 | f | 1 | U | LP | 0 | 0 | 0 | 0 |
| Pat_059 | m | 0 | M | HP | 1 | 0 | 0 | 0 |
| Pat_060 | m | 1 | M | IP | 1 | 0 | 0 | 1 |
| Pat_061 | m | 0 | U | LP | 0 | 0 | 1 | 0 |

*(continued)*

| Patient ID | Sex | Treated before | IGHV status | Methylation Cluster | Del13q | Del11q | Trisomy 12 | Del17p |
|---|---|---|---|---|---|---|---|---|
| Pat_062 | m | 1 | U | LP | 1 | 1 | 0 | 0 |
| Pat_063 | f | 0 | U | LP | 1 | 0 | 0 | 0 |
| Pat_064 | m | 1 | U | LP | 1 | 1 | 0 | 1 |
| Pat_065 | m | 0 | U | LP | 0 | 1 | 0 | 0 |
| Pat_066 | m | 1 | U | LP | 1 | 1 | 0 | 0 |
| Pat_067 | f | 0 | M | HP | 1 | 0 | 0 | 0 |
| Pat_068 | m | 0 | M | HP | 1 | 0 | 0 | 0 |
| Pat_069 | m | 1 | M | HP | 1 | 0 | 0 | 0 |
| Pat_070 | f | 0 | M | HP | 0 | 0 | 0 | 0 |
| Pat_071 | f | 0 | U | LP | 0 | 0 | 0 | 0 |
| Pat_072 | m | 1 | M | HP | 0 | 0 | 1 | 0 |
| Pat_073 | f | 0 | U | LP | 1 | 1 | 0 | 0 |
| Pat_074 | f | 0 | M | HP | 0 | 0 | 0 | 0 |
| Pat_075 | f | 0 | M | HP | 1 | 0 | 0 | 0 |
| Pat_076 | m | 0 | M | HP | 1 | 0 | 0 | 0 |
| Pat_077 | f | 0 | U | NA | 0 | 0 | 1 | 0 |
| Pat_078 | m | 0 | U | LP | 0 | 0 | 0 | 0 |
| Pat_079 | m | 0 | M | HP | 1 | 0 | 0 | 0 |
| Pat_080 | f | 0 | M | HP | 0 | 0 | 0 | 0 |
| Pat_081 | m | 0 | M | HP | 0 | 0 | 0 | 0 |
| Pat_082 | m | 0 | U | LP | 0 | 0 | 0 | 0 |
| Pat_083 | m | 0 | M | HP | 0 | 0 | 0 | 0 |
| Pat_084 | m | 0 | U | LP | 0 | 0 | 1 | 0 |
| Pat_085 | m | 0 | M | HP | 0 | 0 | 0 | 0 |
| Pat_086 | m | 1 | U | LP | 1 | 1 | 0 | 0 |
| Pat_087 | m | 0 | M | IP | 1 | 0 | 0 | 0 |
| Pat_088 | f | 1 | U | LP | 1 | 0 | 0 | 1 |
| Pat_089 | m | 0 | M | HP | 0 | 0 | 0 | 0 |
| Pat_090 | m | 1 | U | LP | 0 | 0 | 1 | 0 |
| Pat_091 | m | 1 | M | NA | 1 | 0 | 0 | 0 |
| Pat_092 | m | 1 | M | HP | 0 | 0 | 0 | 0 |
| Pat_093 | m | 0 | M | HP | 1 | 0 | 0 | 0 |
| Pat_094 | f | 1 | M | HP | 0 | 0 | 0 | 0 |
| Pat_095 | m | 1 | U | LP | 0 | 0 | 0 | 0 |
| Pat_096 | m | 0 | M | HP | 0 | 0 | 0 | 0 |
| Pat_097 | f | 1 | U | LP | 0 | 0 | 0 | 0 |
| Pat_098 | m | 0 | M | HP | 1 | 0 | 0 | 0 |
| Pat_099 | f | 0 | U | LP | 1 | 0 | 0 | 0 |
| Pat_100 | f | 0 | U | LP | 1 | 0 | 0 | 0 |
| Pat_101 | f | 0 | U | LP | 1 | 0 | 0 | 0 |
| Pat_102 | m | 1 | U | IP | 0 | 0 | 0 | 0 |
| Pat_103 | f | 0 | M | HP | 1 | 0 | 1 | 0 |
| Pat_104 | f | 1 | M | HP | 0 | 0 | 0 | 0 |
| Pat_105 | m | 0 | U | LP | NA | NA | NA | NA |
| Pat_106 | m | 0 | U | LP | 1 | 1 | 0 | 1 |
| Pat_107 | f | 0 | M | NA | 1 | 0 | 0 | 0 |
| Pat_108 | m | 0 | M | NA | 1 | 0 | 1 | 0 |
| Pat_109 | m | 0 | M | HP | 1 | 0 | 0 | 0 |
| Pat_110 | m | 0 | M | IP | 1 | 1 | 0 | 0 |
| Pat_111 | f | 1 | U | LP | 1 | 0 | 0 | 0 |
| Pat_112 | f | 0 | M | NA | 0 | 0 | 1 | 0 |
| Pat_113 | f | 0 | M | HP | 1 | 0 | 0 | 0 |
| Pat_114 | m | 0 | U | LP | 1 | 0 | 0 | 0 |
| Pat_115 | m | 1 | U | LP | 1 | 0 | 0 | 1 |
| Pat_116 | m | 0 | U | LP | 1 | 0 | 0 | 0 |
| Pat_117 | m | 0 | U | IP | 1 | 0 | 0 | 0 |
| Pat_118 | f | 0 | M | HP | 0 | 0 | 0 | 1 |
| Pat_119 | m | 0 | U | LP | NA | NA | NA | NA |
| Pat_120 | f | 0 | U | LP | 0 | 0 | 1 | 0 |
| Pat_121 | m | 0 | U | LP | 0 | 0 | 0 | 1 |

*(continued)*

| Patient ID | Sex | Treated before | IGHV status | Methylation Cluster | Del13q | Del11q | Trisomy 12 | Del17p |
|---|---|---|---|---|---|---|---|---|
| Pat_122 | m | 0 | U | LP | 0 | 1 | 0 | 0 |
| Pat_123 | m | 0 | U | LP | 1 | 0 | 0 | 1 |
| Pat_124 | f | 0 | M | HP | 1 | 0 | 1 | 0 |
| Pat_125 | f | 0 | M | HP | 1 | 0 | 0 | 0 |
| Pat_126 | m | 1 | NA | NA | 0 | 0 | 1 | 0 |
| Pat_127 | m | 0 | U | LP | 1 | 1 | 0 | 1 |
| Pat_128 | m | 0 | M | HP | 1 | 0 | 0 | 0 |
| Pat_129 | m | 1 | U | LP | 1 | 0 | 0 | 0 |
| Pat_130 | f | 0 | M | IP | 1 | 0 | 0 | 0 |
| Pat_131 | f | 0 | U | LP | 1 | 0 | 0 | 1 |
| Pat_132 | m | 1 | U | LP | 1 | 0 | 0 | 1 |
| Pat_133 | m | 1 | U | LP | 1 | 0 | 0 | 1 |
| Pat_134 | m | 1 | M | HP | 1 | 0 | 0 | 1 |
| Pat_135 | m | 0 | M | HP | 1 | 0 | 0 | 0 |
| Pat_136 | m | 0 | M | HP | 1 | 0 | 0 | 0 |
| Pat_137 | f | 0 | U | LP | 0 | 0 | 1 | 0 |
| Pat_138 | f | 0 | M | HP | 1 | 0 | 0 | 0 |
| Pat_139 | f | 0 | M | HP | NA | NA | NA | NA |
| Pat_140 | f | 0 | M | IP | 1 | 0 | 0 | 0 |
| Pat_141 | m | 0 | M | IP | 1 | 0 | 0 | 0 |
| Pat_142 | m | 0 | M | HP | 1 | 0 | 0 | 0 |
| Pat_143 | f | 0 | U | LP | 1 | 0 | 0 | 0 |
| Pat_144 | m | 1 | U | LP | 0 | 0 | 0 | 1 |
| Pat_145 | f | 0 | M | HP | 0 | 0 | 0 | 0 |
| Pat_146 | m | 1 | U | LP | NA | NA | NA | NA |
| Pat_147 | m | 1 | U | LP | 0 | 0 | 0 | 1 |
| Pat_148 | m | 1 | U | NA | 1 | 0 | 0 | 0 |
| Pat_149 | m | 1 | U | LP | 0 | 1 | 1 | 0 |
| Pat_150 | f | 0 | M | HP | 1 | 0 | 0 | NA |
| Pat_151 | m | 0 | U | LP | 0 | 0 | 1 | 0 |
| Pat_152 | m | 0 | NA | NA | 0 | 1 | 1 | 0 |
| Pat_153 | m | 1 | U | LP | 1 | 0 | 0 | 0 |
| Pat_154 | m | 0 | M | HP | 1 | 0 | 0 | 0 |
| Pat_155 | m | 0 | M | HP | 1 | 0 | 0 | 0 |
| Pat_156 | f | 0 | U | LP | 1 | 0 | 0 | 0 |
| Pat_157 | f | 0 | U | LP | 1 | 1 | 0 | 0 |
| Pat_158 | m | 0 | M | HP | 1 | 0 | 0 | 0 |
| Pat_159 | f | 0 | U | LP | 1 | 0 | 0 | 0 |
| Pat_160 | m | 0 | U | LP | NA | NA | NA | NA |
| Pat_161 | m | 1 | U | LP | 0 | 0 | 0 | 0 |
| Pat_162 | m | 0 | M | HP | 0 | 0 | 0 | 0 |
| Pat_163 | m | 0 | M | HP | 0 | 0 | 0 | 0 |
| Pat_164 | f | 0 | U | LP | 0 | 0 | 0 | 0 |
| Pat_165 | m | 1 | U | LP | 0 | 1 | 1 | 0 |
| Pat_166 | m | 0 | U | LP | 1 | 0 | 1 | 0 |
| Pat_167 | m | 0 | NA | IP | NA | NA | NA | NA |
| Pat_168 | m | 0 | M | HP | 1 | 1 | 0 | 0 |
| Pat_169 | m | 0 | M | HP | 0 | 0 | 1 | 0 |
| Pat_170 | m | 0 | M | HP | 1 | 0 | 0 | 0 |
| Pat_171 | m | 0 | M | HP | 1 | 0 | 0 | 0 |
| Pat_172 | m | 0 | M | HP | 1 | 0 | 0 | 0 |
| Pat_173 | m | 0 | M | HP | 1 | 0 | 0 | 0 |
| Pat_174 | f | 0 | U | LP | NA | 0 | NA | 0 |
| Pat_175 | m | 0 | U | LP | 0 | 0 | 0 | 0 |
| Pat_176 | f | 0 | M | HP | NA | NA | NA | NA |
| Pat_177 | f | 0 | NA | IP | NA | 1 | NA | NA |
| Pat_178 | m | 0 | M | HP | 1 | 0 | 0 | 0 |
| Pat_179 | m | 0 | U | LP | 0 | 1 | 0 | 0 |
| Pat_180 | m | 0 | M | HP | NA | NA | NA | NA |
| Pat_181 | f | 0 | M | HP | NA | NA | NA | NA |

*(continued)*

| Patient ID | Sex | Treated before | IGHV status | Methylation Cluster | Del13q | Del11q | Trisomy 12 | Del17p |
|---|---|---|---|---|---|---|---|---|
| Pat_182 | m | 0 | M | HP | 1 | 0 | 0 | 0 |
| Pat_183 | f | 1 | NA | IP | 1 | 0 | 0 | 0 |
| Pat_184 | m | 0 | U | LP | NA | NA | NA | NA |
| Pat_185 | m | 0 | M | HP | NA | NA | NA | NA |
| Pat_186 | m | 0 | M | HP | 1 | 0 | 0 | 0 |
| Pat_187 | m | 1 | U | LP | NA | NA | NA | NA |
| Pat_188 | m | 0 | NA | NA | NA | NA | NA | NA |
| Pat_189 | m | 0 | NA | NA | NA | NA | NA | NA |
| Pat_190 | m | 0 | NA | NA | NA | NA | NA | NA |
| Pat_191 | f | 0 | NA | NA | 0 | 0 | 0 | 0 |
| Pat_192 | f | 0 | NA | NA | 1 | 0 | 0 | 0 |

**Supplementary Table 3. Patient samples included in the study.** List of patient samples and selected characteristics. For a full list of characteristics see online vignette.

| Factor | HR | p value | CI Low | CI High |
|---|---|---|---|---|
| Cluster 3 vs Cluster 1 | 0.96 | 0.89 | 0.54 | 1.72 |
| Cluster 3 vs Cluster 2 | 1.68 | 0.17 | 0.80 | 3.51 |
| Cluster 3 vs Cluster 4 | 0.44 | 0.04 | 0.20 | 0.96 |
| IGHV.status | 1.74 | 0.04 | 1.02 | 2.96 |
| trisomy 12 | 0.87 | 0.71 | 0.44 | 1.76 |
| TP53 | 4.01 | <0.0001 | 2.41 | 6.69 |

**Supplementary Table 4. Multivariate survival analysis of response clusters.** Multivariate Cox proportional hazards model of stimuli response clusters and genetic subgroups of disease progression using TTT and C3 as reference.

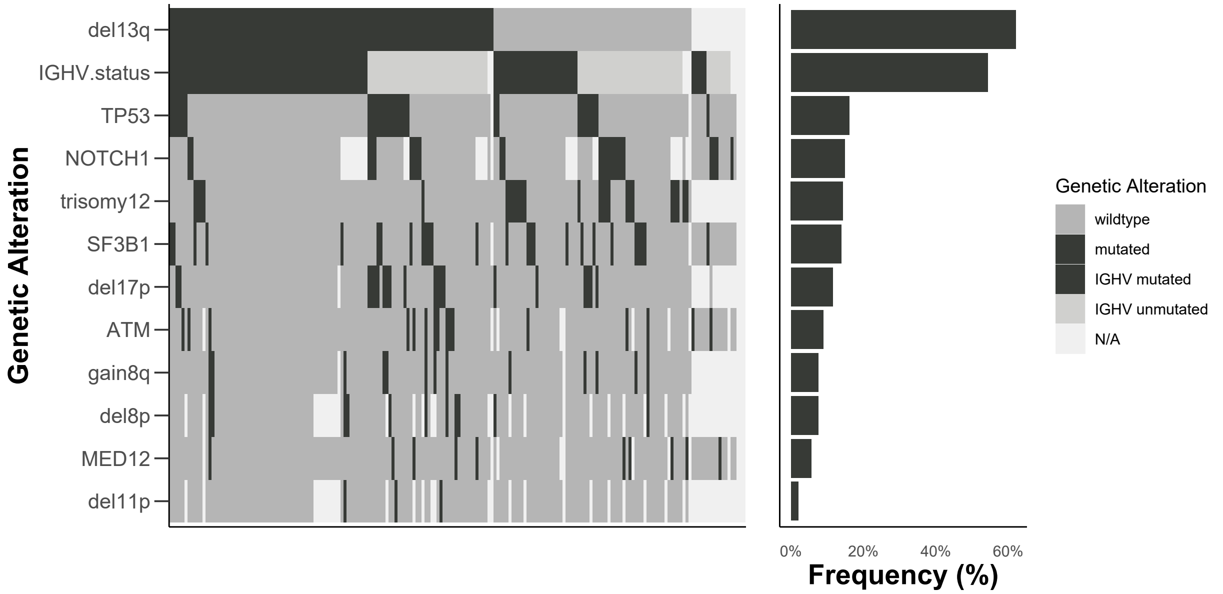


**Supplementary Figure 1. Genetic profiles of screened patient samples.** Selected genetic alterations on y-axis and screened patient samples on x-axis.

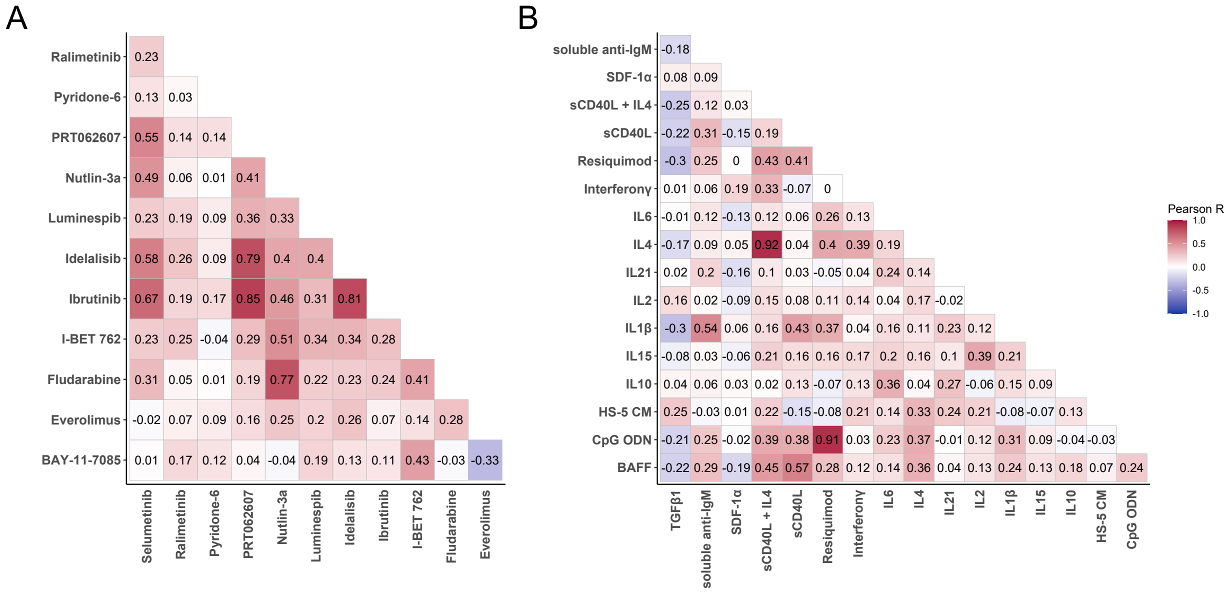


**Supplementary Figure 2. Correlation of responses to stimuli and drugs.** Pearson correlation of log transformed viability values normalised to untreated control. Correlation between drug treatments **(A)** and microenvironmental stimuli **(B)**.

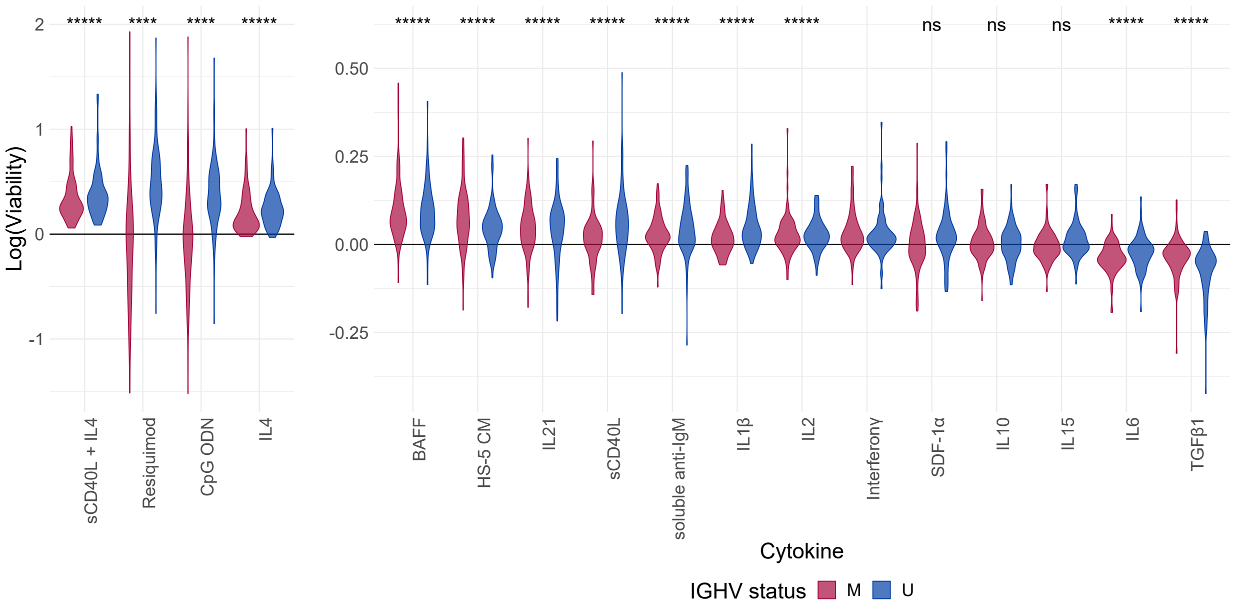


**Supplementary Figure 3. Response to stimuli by IGHV status.** Viability after 48h incubation with microenvironmental stimuli. Log transformed viabilities, normalized to DMSO solvent controls, stratified by IGHV status. BH-adjusted p-values are shown from one-sample t-tests of all patient samples. (p<0.00001=*****, p<0.0001=****, p>0.05=ns)

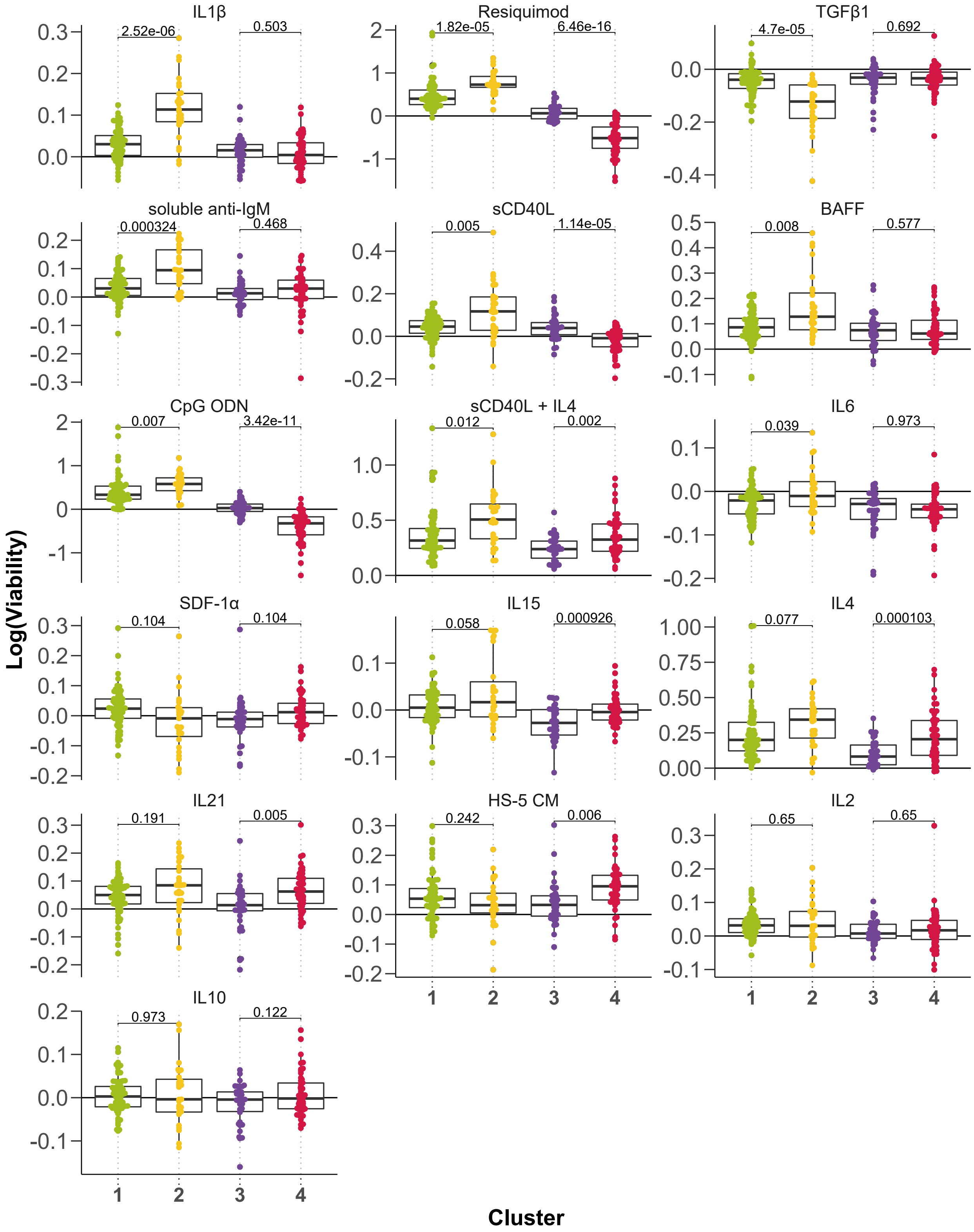


**Supplementary Figure 4. Stimuli response by clusters.** Log transformed viabilities after treatment with stimuli, faceted by cluster. BH-adjusted p-values from student's t-tests are shown.

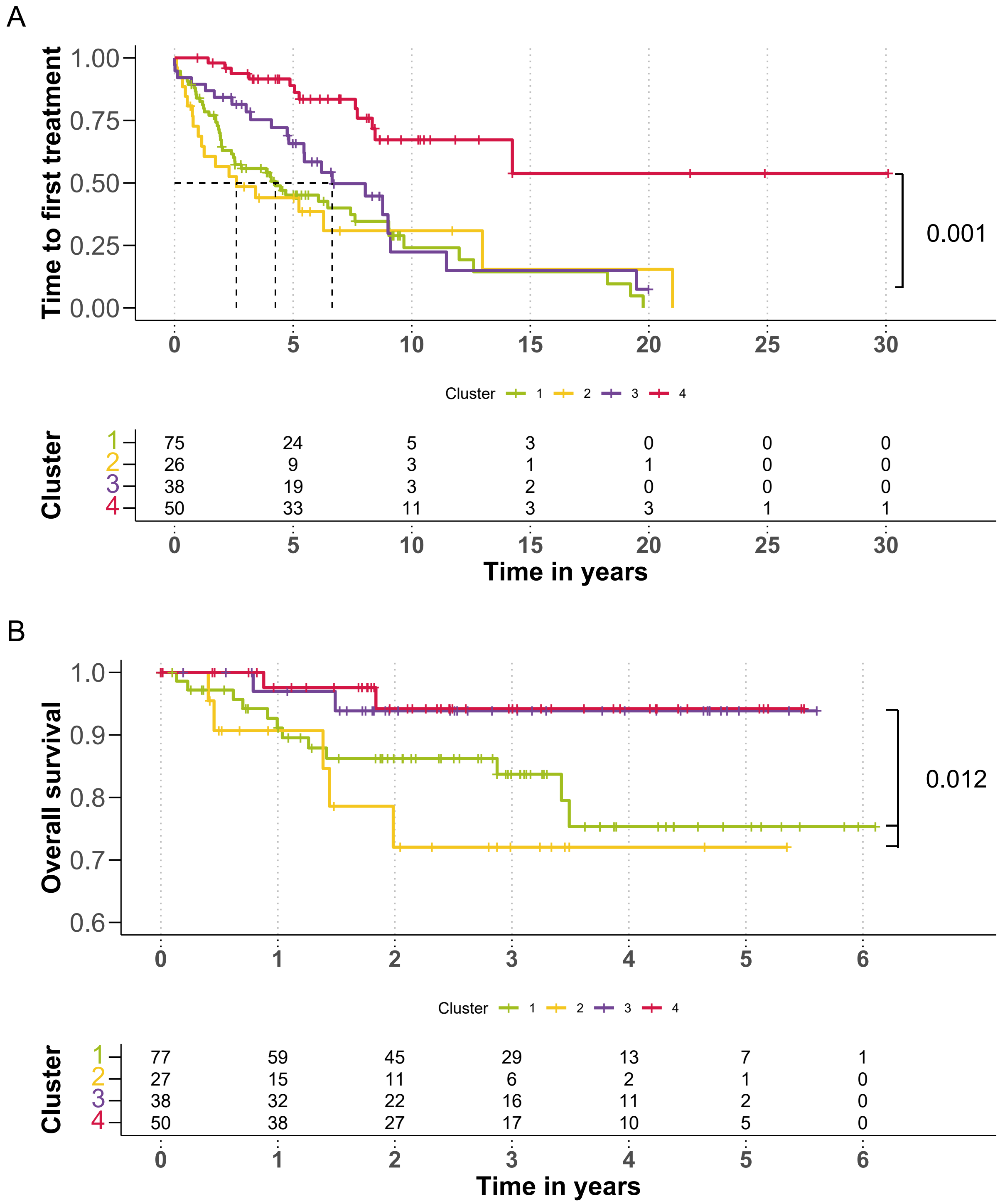


**Supplementary Figure 5. Time to first treatment and overall survival by clusters.** Kaplan-Meier Curves of time to first treatment with p-value from univariate Cox proportional hazards model between C3 and C4 **(A)** and overall survival with p-value from univariate Cox proportional hazards model between C1&2 and C3&4 **(B)**. Median survival not reached.

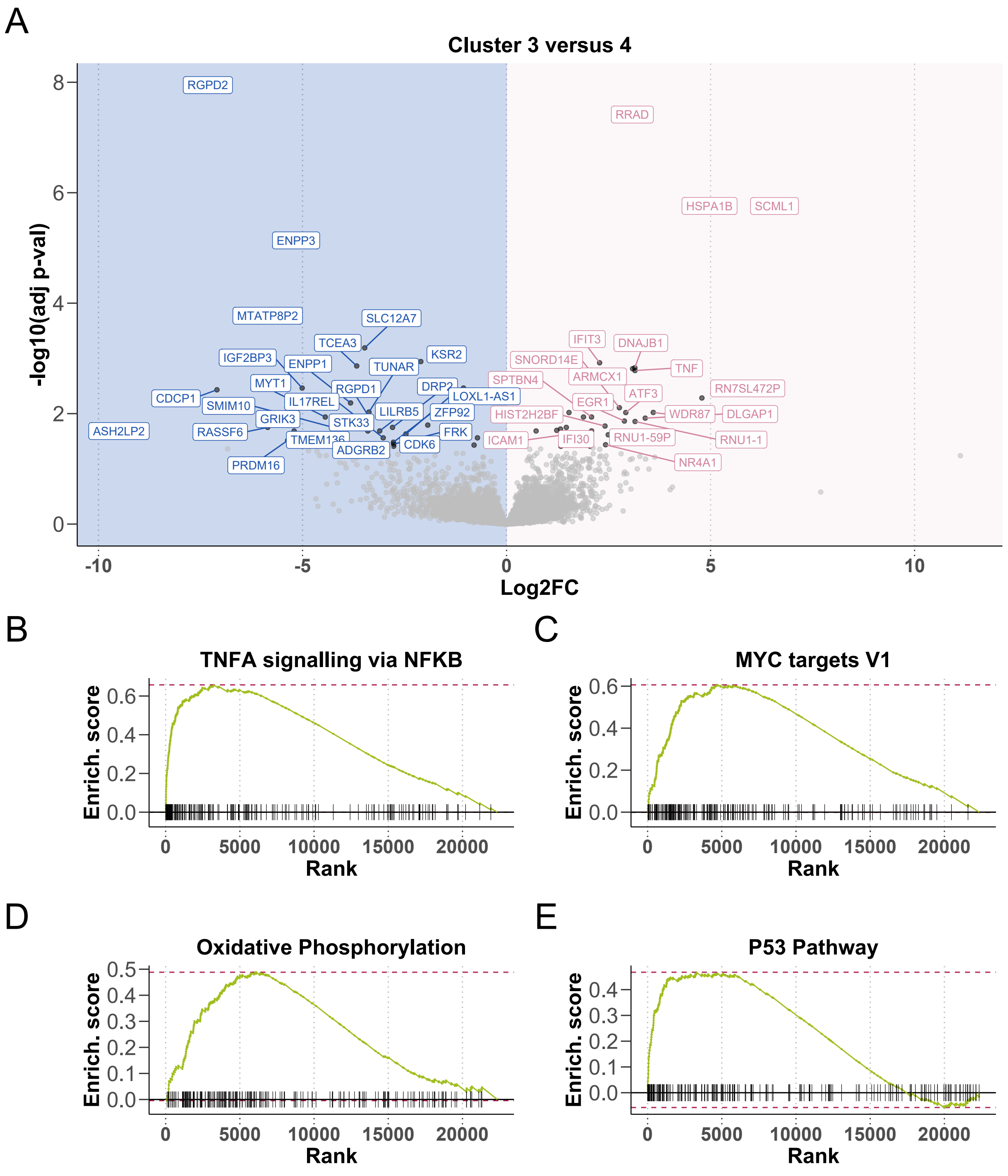


**Supplementary Figure 6. RNA-Sequencing of matched samples indicates differential gene expression between Cluster 3 and Cluster 4.** Volcano plot of differentially expressed genes between Cluster 3 and Cluster 4 **(A)**. X axis indicates log2 fold change values, calculated using the Deseq package, y axis gives corresponding -log10(adjusted p value). P values adjusted using BH method. Genes are labeled where p value < 0.05. Enrichment plots of selected pathways **(B-D)**. Gene set enrichment analysis (GSEA) was performed with the Hallmark gene sets from the GSEA Molecular Signatures Database. Wald statistic was used to rank the genes. The green curve corresponds to the Enrichment Score curve, which is the running sum of the weighted enrichment score obtained from GSEA software.

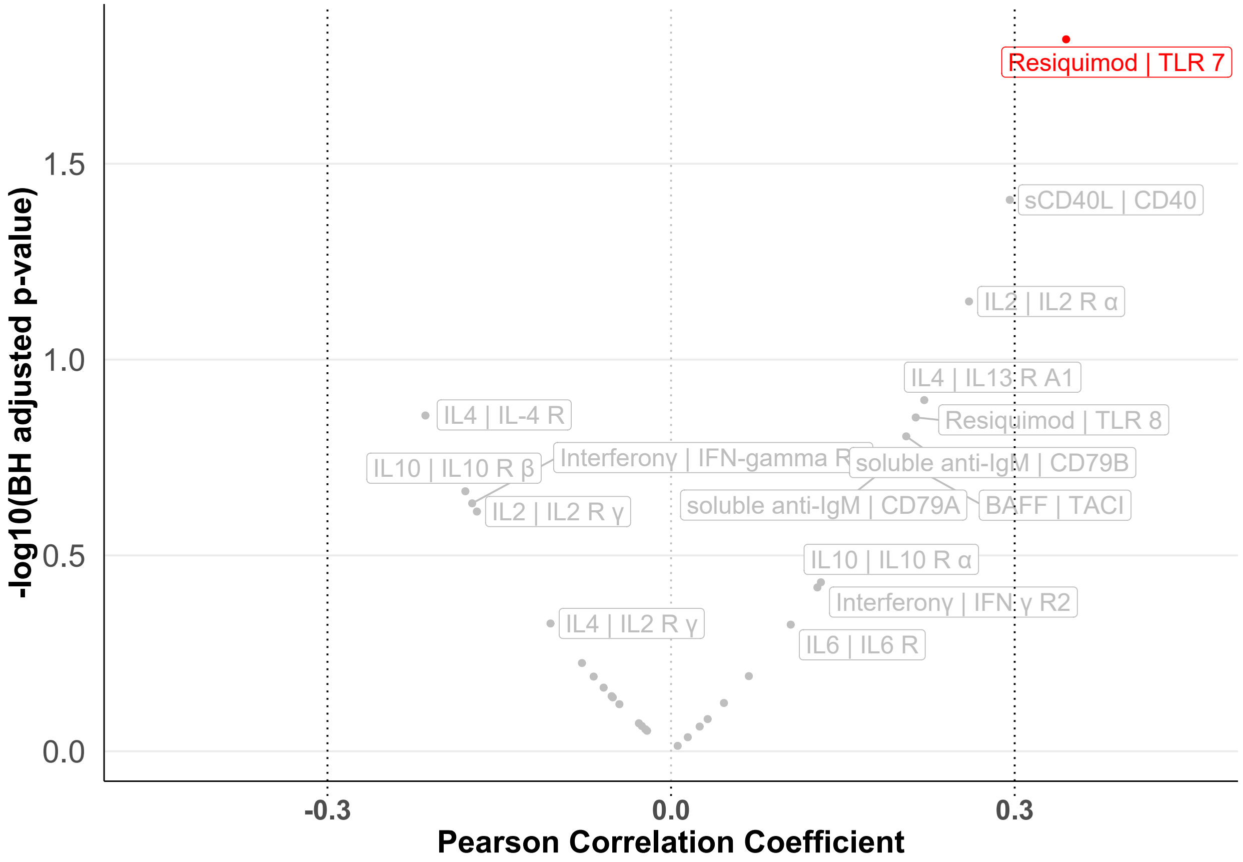


**Supplementary Figure 7. Correlation of stimuli response and receptor expression.** The effects of the microenvironmental stimuli on viability were largely independent of the expression of the corresponding stimuli receptors. Volcano plot depicts Pearson correlation coefficients against corresponding BH-adjusted p values, for the correlation between control - normalised log viability values with each stimulus and vst RNA counts of corresponding stimuli receptor. RNA counts taken from RNA-Sequencing of untreated matched CLL patient sample. Only viability after treatment with Resiquimod correlated with receptor expression (R>0.3).

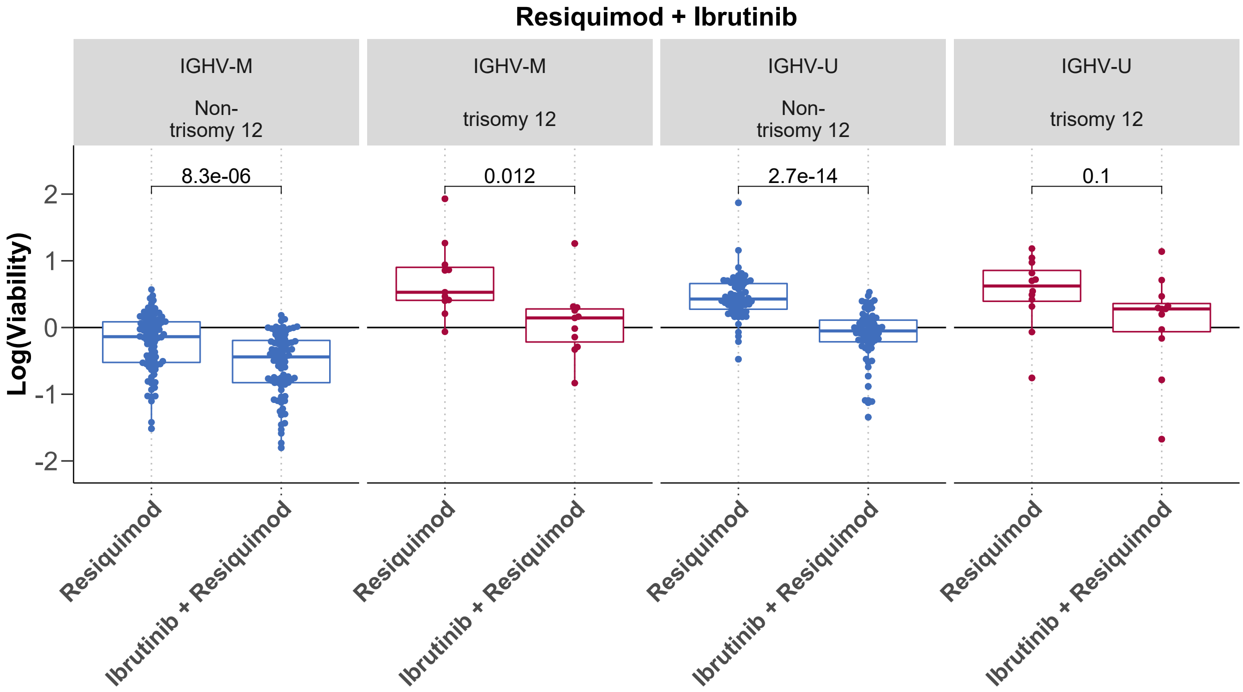


**Supplementary Figure 8. BCR inhibition by ibrutinib counters the protective effect of TLR stimulation in all genetic subgroups of IGHV and trisomy 12.** Log normalized viability after treatment with resiquimod with and without ibrutinib. Faceted by IGHV mutation status and trisomy 12.

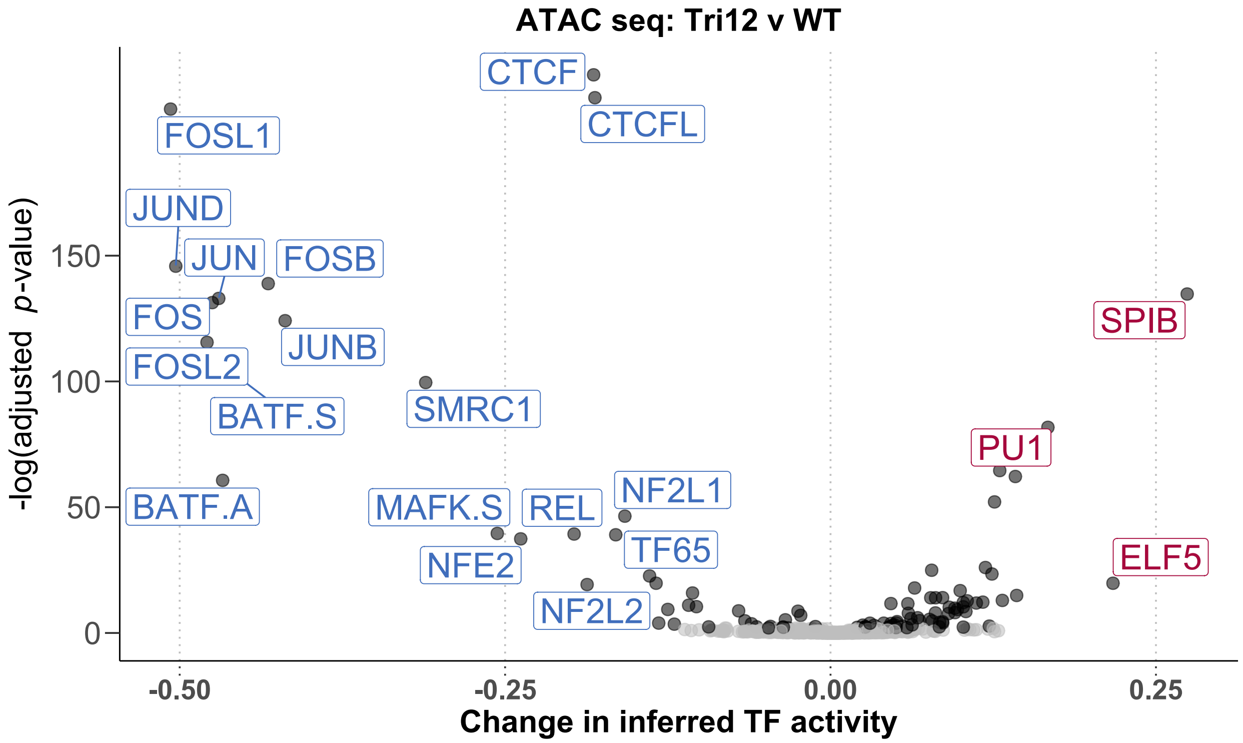


**Supplementary Figure 9. ATAC-Seq comparing trisomy 12 and non-trisomy 12 CLL samples.** Volcano plot shows change in inferred TF activity (x axis) against adjusted p-values (y axis) for trisomy 12 (n = 2) versus non-trisomy 12 samples (n = 2). Inferred TF activity calculated using the diffTF package, and measured as weighted mean difference. p-values are obtained through diffTF in analytic mode and adjusted by the Benjamini-Hochberg procedure. TFs are labeled where absolute weighted mean difference > 0.15, and adjusted p-value < 0.01.

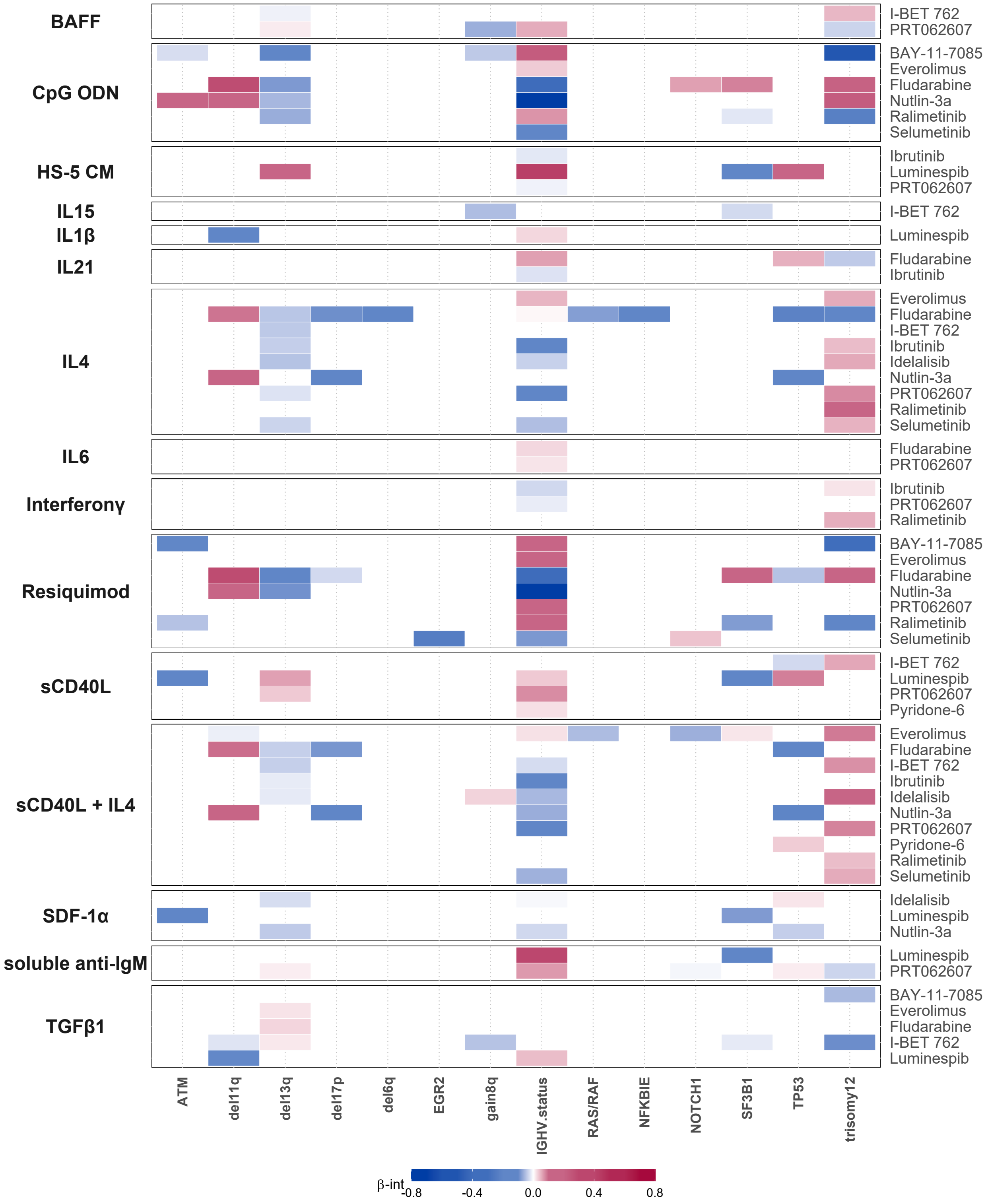


**Supplementary Figure 10. Genetic predictors of drug - stimulus interactions.** Heatmap depicting overview of genetic predictors of drug - stimulus interactions (each row represents the coefficients from fitting a single multivariate model). Stimuli are shown on left, and corresponding drugs on right. Drugs, stimuli and genetic alterations are alphabetically sorted. Coloured fields indicate that the $\beta_{int}$ for given drug and stimulus is modulated by corresponding genetic feature. Positive coefficients are shown in red, indicating $\beta_{int}$ is more positive for given drug and stimulus combination if the feature is present.

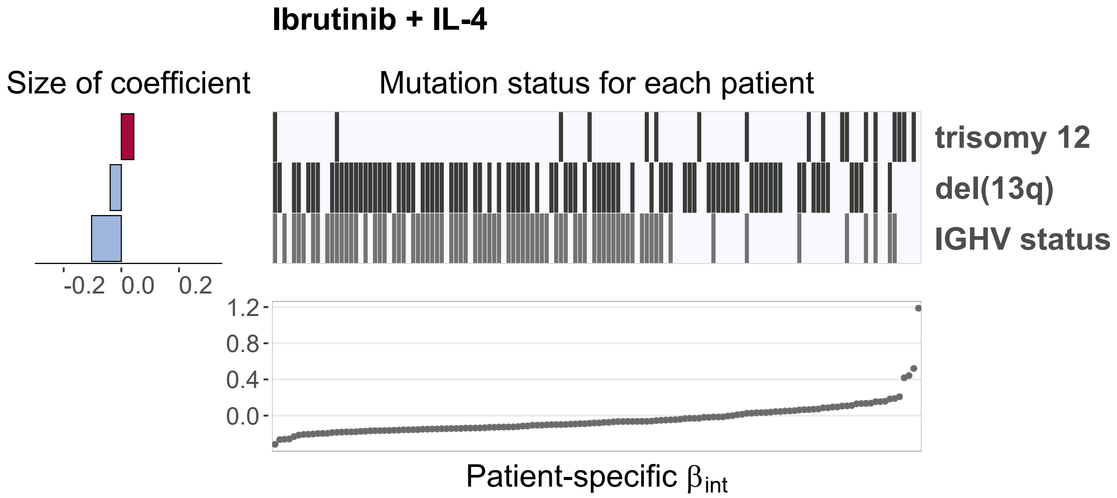


**Supplementary Figure 11. Genetic predictors of the interaction between ibrutinib and IL4.** Predictor profile depicting genetic features that modulate the size of $\beta_{int}$ for ibrutinib and IL4. To generate predictor profile, linear model in Eqn. (1) was fitted in a sample - specific manner, to calculate drug-stimulus interaction coefficients ($\beta_{int}$) for each patient sample. Ranked patient-specific $\beta_{int}$ values are shown in lower scatter plot. Associations between the size of $\beta_{int}$ and genetic features were identified using multivariate regression with L1 (lasso) regularisation, with gene mutations and IGHV status as predictors, and selecting coefficients that were chosen in >90% of bootstrapped model fits. The horizontal bars on left show the size of fitted coefficients assigned to genetic features. Matrix above scatter plot indicates patient mutation status for the selected genetic features. Matrix fields correspond to points in scatter plot (ie patient data is aligned), to indicate how the size of $\beta_{int}$ varies with selected genetic feature.

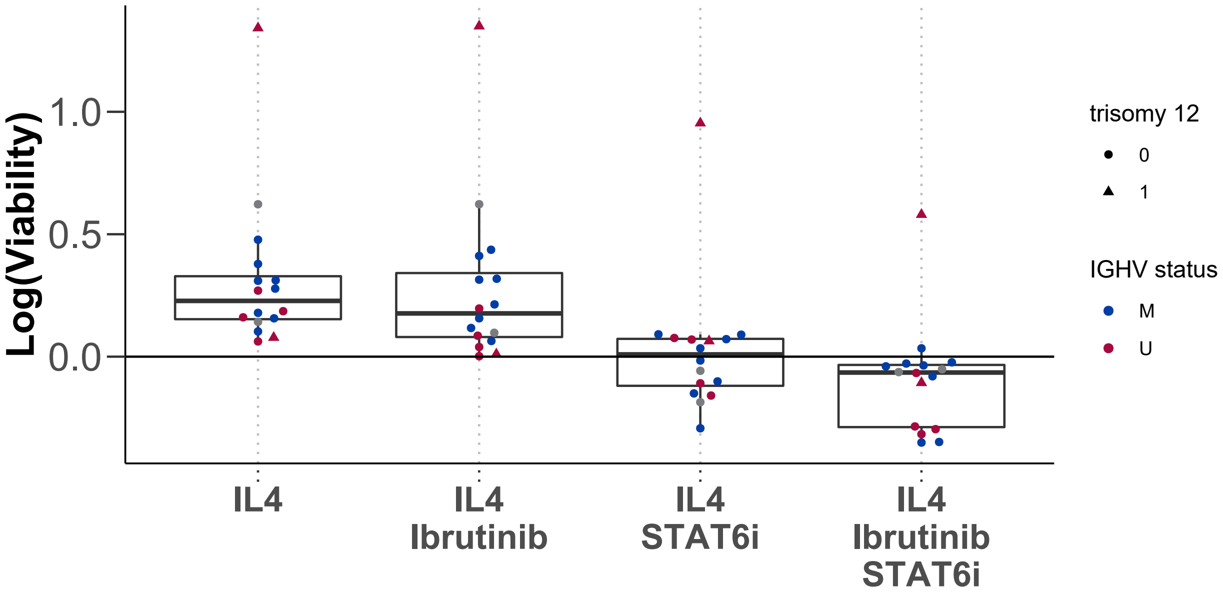


**Supplementary Figure 12. STAT6 dependency of IL4 signaling.** The effects observed with IL4 stimulation are dependent on STAT6 activation. Addition of the STAT6 inhibitor AS1517499 could revoke the effect on baseline viability as well as drug induced toxicity. Log transformed viability after treatment with IL4 in combination with ibrutinib, the STAT6 inhibitor AS1517499, and both.

### References


1. Dietrich S, Oleś M, Lu J, et al. Drug-perturbation-based stratification of blood cancer. J. Clin. Invest. 2018;128(1):427–445.

2. Buenrostro JD, Wu B, Chang HY, Greenleaf WJ. ATAC-seq: A Method for Assaying Chromatin Accessibility Genome-Wide. Curr. Protoc. Mol. Biol. 2015;109:21.29.1–21.29.9.

3. Berest I, Arnold C, Reyes-Palomares A, et al. Quantification of Differential Transcription Factor Activity and Multiomics-Based Classification into Activators and Repressors: diffTF. Cell Rep. 2019;29(10):3147–3159.e12.

4. Kulakovskiy IV, Vorontsov IE, Yevshin IS, et al. HOCOMOCO: expansion and enhancement of the collection of transcription factor binding sites models. Nucleic Acids Res. 2016;44(D1):D116–25.
